# A role for CASM in the repair of damaged Golgi architecture

**DOI:** 10.1101/2025.09.04.674289

**Authors:** Seeun Oh, Saif Ullah, Bhaskar Saha, Michael A Mandell

## Abstract

The term CASM describes a process in which MAP1LC3B/LC3B and other Atg8-family proteins are covalently ligated to lipids in damaged endomembranes. While CASM is commonly described as a cytoprotective response to multiple types of membrane damage, how CASM helps cells maintain homeostasis is still unclear. Here, we show that CASM maintains Golgi apparatus architecture following the loss of TRIM46, a ubiquitin ligase with roles in microtubule organization. TRIM46 deficient cells were notable for enhanced TFEB-driven lysosomal biogenesis and Golgi ribbon fragmentation, with colocalization of the *trans*-Golgi marker TGOLN2 and the Atg8-family proteins LC3B and GABARAP. Further studies revealed that the Golgi Atg8ylation seen in *TRIM46* knockout cells was not degradative and mechanistically resembled CASM. Genetic inhibition of CASM in TRIM46 deficient cells reduced TFEB activation and exacerbated the Golgi morphology defects, suggesting that CASM contributes to Golgi repair. Accordingly, Golgi reformation after drug-induced fragmentation was impaired upon knockdown of CASM genes. Together, these studies identify lysosomal biogenesis and CASM as coordinated features of a Golgi damage response, with CASM acting to preserve Golgi integrity.

## Introduction

The proper organization, placement, and maintenance of membranous organelles is fundamental to the life of eukaryotic cells. This requires the repair or replacement of damaged membranes or dysfunctional membrane-associated proteins. Multiple mechanisms of membrane repair have evolved, likely due to the critical importance of maintaining membrane integrity and function, the diversity of organelle membranes within a cell, and the variety of possible causes of membrane damage. Depending on the type of membrane damage, cellular responses can include: i) ESCRT-mediated excision of damaged membranes; ii) delivery of new membrane to the site of damage by organelle fusion or by membrane-to-membrane lipid transfer; and iii) enhanced expression and biosynthesis of protein constituents of damaged membranes [1]. Additionally, multiple membrane damage scenarios trigger the attachment of a set of ubiquitin-like proteins, collectively referred to as Atg8-family proteins (LC3A, LC3B, LC3C, GABARAP, GABARAPL1, and GABARAPL2), to membranes in a process termed “Atg8ylation” [2–4].

Membrane Atg8ylation contributes to membrane damage responses in several ways. These include enabling the lysosome-mediated removal of damaged membranes or membranous organelles [5–8], indirectly up-regulating the biosynthesis of new membrane components [9–11], and providing a “patch” to plug damaged membranes [12]. Mechanistically, the Atg8ylation process resembles ubiquitylation, with the roles of E1 activating enzyme and E2 conjugating enzyme played by ATG7 and ATG3, respectively [3]. There are two multi-protein E3 ligase complexes that are known to act in Atg8ylation: the ATG12–ATG5-ATG16L1 complex [13] and the ATG12–ATG5-TECPR1 complex [14–16]. The E3 ligase complexes attach Atg8 proteins to either phosphatidylethanolamine (PE) or phosphatidylserine (PS). Macroautophagy (hereafter, autophagy) is the best-studied pathway that utilizes membrane Atg8ylation. In autophagy, Atg8 proteins are attached to *de novo*-created membrane that elongates in a curved manner and ultimately sequesters cytoplasmic contents within a sealed double-membraned vesicle termed an autophagosome [17]. Autophagosomes are then trafficked along microtubules towards lysosomes, where the two organelles fuse and the inner autophagosomal membrane and its luminal contents are degraded [18]. More recently, additional processes that require the Atg8ylation machinery have been identified. These processes differ from autophagy in that they involve Atg8ylation of pre-existing membranes, they do not result in the formation of autophagosomes, and they are not always degradative [2]. The term <u>c</u>onjugation of <u>A</u>tg8 to <u>s</u>ingle <u>m</u>embranes (CASM) has been coined to describe these Atg8ylation-dependent pathways [19]. While both autophagy and CASM utilize the same Atg8ylation machinery, they differ in the mechanism by which the E3 ligase complex is recruited to the sites of Atg8ylation. In autophagy, the ATG16L1-containing E3 complex is recruited to phosphatidylinositol-3-phosphate containing membranes generated by the BECN1 (beclin 1)-PIK3C3/VPS34 complex. The BECN1 complex is activated by a protein complex containing the kinase ULK1 or ULK2. On the other hand, BECN1 and the ULK1 complex are dispensable for CASM [19–21]. Finally, the two pathways differ in terms of which lipid is conjugated to Atg8-family proteins, with PE being exclusively utilized by autophagy whereas both PE and PS are utilized by CASM [22]. Despite their mechanistic differences, both autophagy and CASM are typically considered protective responses to membrane damage, although the functions of CASM remain largely undefined.

Ubiquitination can be a key regulator of Atg8ylation-dependent processes [23]. This is particularly true in the case of autophagy. For example, many members of the tripartite motif containing (TRIM) family of ubiquitin ligases are reported to act as both autophagy regulators and in the identification of membranous or proteinaceous autophagy substrates [24–26]. The role(s) of TRIM proteins, or ubiquitination more generally, in other Atg8ylation-dependent processes such as CASM have not yet been explored.

Here, our investigation into the actions of TRIM proteins in Atg8ylation led us to find that disruption of microtubule organization is a potent activator of the non-degradative Atg8ylation of *trans*-Golgi membranes. This subsequently activates a program of lysosomal biogenesis and autophagy. We found that TRIM46, a ubiquitin ligase previously implicated in the formation of an axonal structure in neurons [27], is important for microtubule organization. Genetic depletion of *TRIM46* resulted in fragmentation of Golgi ribbon structure and activation of TFEB and TFE3, the master transcriptional activators of lysosome- and autophagy-related gene expression. *TRIM46* knockout also resulted in substantial Atg8ylation of Golgi membranes in a manner reminiscent of CASM. When Atg8ylation was blocked, the Golgi morphology defects seen in *TRIM46* knockout cells were exacerbated, while the activation of TFEB was attenuated. The activation of both CASM and TFEB seen in *TRIM46* knockout is phenocopied by chemical inhibitors of microtubule function or vesicle trafficking, all of which disrupt Golgi architecture. Importantly, the Atg8ylation machinery was required for the reestablishment of normal Golgi morphology following chemical-mediated Golgi fragmentation. Overall, our study indicates that CASM-dependent activation of lysosomal biogenesis is a general response to perturbations of Golgi architecture, with CASM having a role in membrane reorganization.

## Results

### Screening of TRIM proteins for roles in autophagosome maturation

Since TRIM proteins have many reported roles in regulating autophagy initiation [24,28,29], we wondered if their actions in autophagy extend to mediating autophagosome-lysosome fusion and autophagy flux. To address this question, we reanalyzed data from a previous siRNA screen that examined the effect of TRIM knockdown on the abundance of the autophagosome marker LC3B [30]. This screen made use of cells expressing mCherry-eYFP-LC3B (tandem fluorescent LC3B, tfLC3B) as a marker of autophagosomes. Although tfLC3B can be used to monitor both the formation of autophagosomes and their delivery to lysosomes (termed autophagosome maturation) due to the differential sensitivity of the two fluorophores to low pH [31,32], in our initial analysis we only considered the abundance of YFP-positive LC3B structures [30]. Here, we determined whether the siRNA-mediated knockdown of any one of 68 TRIMs in HeLa cells altered the delivery of the LC3B reporter to acidified compartments, which can be detected by comparing the relative abundance of “non-acidified” LC3B structures (positive eYFP signal) with that of “total” LC3B structures (positive for mCherry signal; Fig. 1A, B and S1A). We found that cells subjected to knockdown of six TRIMs (TRIMs 25, 38, 46, 47, 61, and 68) reduced the acidification of the LC3B reporter by more than three standard deviations below the mean of cells transfected with non-targeting siRNAs in two out of two experiments (Fig. 1B and S1A). This indicates a possible role for these TRIMs in autophagosome maturation. Alternatively, these TRIMs could act to inhibit non-degradative Atg8ylation pathways such as CASM.

**Figure 1.**
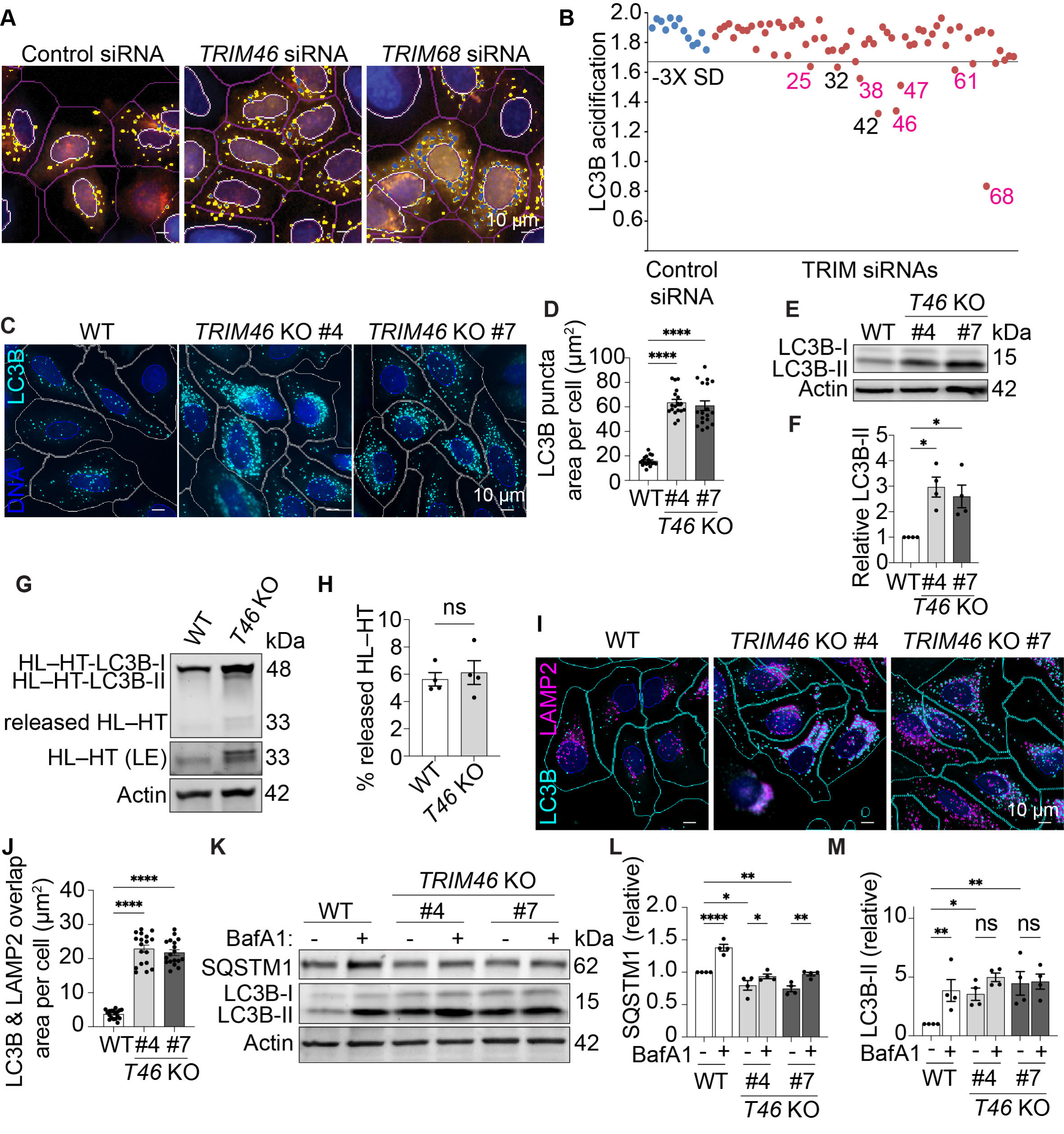
TRIM46 regulates nondegradative LC3 lipidation. (**A**) siRNA screen for TRIMs regulating the accumulation of non-acidified LC3B. HeLa cells stably expressing mCherry-eYFP-LC3B were transfected with TRIM or control siRNA two days prior to fixation and high content imaging. Neutral pH LC3B puncta (eYFP positive; blue mask) or total LC3B puncta (yellow mask) were identified and quantitated from >500 cells per siRNA. White mask, nuclei; magenta mask, cell boundary. (**B**) TRIM knockdowns (red data points) with LC3B acidification reduced by more than three standard deviations (3X SD) below the mean of non-targeting siRNA controls (blue data points) were identified as hits. Magenta numbers indicate TRIMs ‘hits’ that were identified in two out of two experiments (see also Figure S1A). (**C–D**) High-content imaging analysis of LC3B abundance in WT and two independent *TRIM46* knockout clones. Each data point represents the average LC3B puncta area from >500 cells. (**E, F**) Immunoblot analysis of LC3B-II levels in *TRIM46* knockout cells. Quantification of LC3B-II from 3 independent experiments. (**G, H**) Halo-LC3 assay for autophagy flux. WT and *TRIM46* knockout cells stably expressing HT-LC3B were pulsed with TMR-HL for 30 min prior to 6-h chase with full media. HT/HL were detected by in-gel fluorescence. LE, long exposure. Quantification (H) shows the percent of total HT/HL signal that results from the “released” HT/HL, a product of lysosomal degradation. (**I, J**) High-content imaging analysis of LC3B (cyan) and LAMP2 (magenta) localization in WT and two *TRIM46* knockout clones. Each data point in (J) represents the average area of LC3B and LAMP2 overlap per cell from >500 cells. (**K-M**) Immunoblot analysis of LC3B-II and SQSTM1 abundance in lysates from WT or *TRIM46* knockout HeLa cells treated with DMSO or 100 nM bafilomycin A_1_ (BafA1) for 6 h. Plots show quantification of relative protein abundance of SQSTM1 (L) and LC3B-II (M). Each dot represents an independent experiment. Statistical analyses were performed using an unpaired t-test (H), ordinary one-way ANOVA followed by Dunnett’s multiple-comparison test (D, F, J), or two-way ANOVA followed by Tukey’s multiple-comparison test (L, M). Data: mean ± SEM; *, p < 0.05; ***, p < 0.001; ****, p < 0.0001; ns, not significant.

### Knockdown of TRIM46 increases non-degradative Atg8ylation without impairing autophagy flux

We chose to focus on TRIM46 as a possible regulator of autophagosome maturation because of its reported actions on microtubule organization in neurons [27], which we reasoned could impact proper autophagosome and/or lysosome trafficking. We generated several clonal populations of HeLa and HEK293T cells in which *TRIM46* was knocked out by CRISPR/Cas9 (Fig. S1B, C). High content imaging of two of the HeLa cell lines showed that the abundance of endogenous LC3B puncta was elevated by ∼3-fold in the *TRIM46* knockout cells relative to wild-type HeLa cells stably expressing Cas9 and non-targeting gRNA (Fig. 1C, D). Immunoblot experiments also showed that *TRIM46* knockout HeLa cells (Fig. 1E, F) or HEK293T cells (Fig. S1D) showed strongly elevated levels of LC3B-II, the form of LC3B that is generated by Atg8ylation. Exogenous expression of TRIM46 reversed the effect of *TRIM46* knockout on LC3-II abundance (Fig. S1E, F). Together, these data show that TRIM46 attenuates the accumulation of lipidated LC3B.

Although the purpose of our initial screen was to identify TRIMs that might be required for autophagosome maturation, further study showed that *TRIM46* deficiency did not impair autophagy flux. This was first demonstrated using the Halo-LC3 assay, which relies on the fact that Halo ligand–Halo tag adducts (HL–HT) are protected from lysosomal degradation. When autophagy flux is active, the lysosome will incompletely degrade HL–HT-fused LC3 (48 kDa), leaving a 33 kDa HL–HT fragment that can readily be detected in SDS-PAGE gels when the Halo ligand is fused to a fluorescent molecule [33]. We detected the HL–HT fragment in lysates from both WT and *TRIM46* knockout cells expressing HT-LC3B and treated with tetramethylrhodamine (TMR)-labeled HL under basal autophagy conditions (Fig. 1G). Although more released HL–HT is present in *TRIM46* knockout cells, quantitation of the released HL–HT band relative to the total abundance of TMR signal (Fig. 1H) indicated that WT and *TRIM46* knockout cells had comparable levels of autophagy flux. Image analysis showed that colocalization between LC3B and the lysosome marker LAMP2 in both lines of *TRIM46* knockout HeLa cells was elevated relative to WT (Fig. 1I, J and S1G, H), indicating no impairment in autophagosome/lysosome fusion. Treatment with the lysosomal inhibitor Bafilomycin A1 (BafA1) increased the abundance of the autophagy substrate SQSTM1 in both WT and TRIM46 knockout cells (Fig. 1K, L), demonstrating functional autophagy flux.

Interestingly, however, the effect of BafA1 on LC3-II levels in *TRIM46* knockout cells was muted relative to what was measured in WT cells in which it strongly increased the abundance of LC3-II (Fig. 1K, M). Similar results were seen when using the Halo-LC3 assay (Fig. S1I). BafA1 treatment increased the abundance of HL–HT-LC3B-II in WT cells pulsed with TMR-HL by ∼3-fold. In contrast, BafA1 had no effect on the abundance of HL–HT-LC3B-II in the *TRIM46* knockout cells. As discussed later, our interpretation for this result as well as those from our tfLC3B experiments (Fig. 1B and S1A) is that not all of the membrane-associated LC3B in *TRIM46* knockout cells is delivered to acidified compartments for degradation and that Atg8ylation pathways in addition to autophagy are activated in TRIM46-deficient cells.

### TRIM46 restrains lysosomal biogenesis

In the experiments shown in Figure 1I and S1E, we noted that the number of LAMP2-positive structures appeared elevated in the *TRIM46* knockout cells, a result that we confirmed by high content imaging (Fig. 2A, B). The expression of lysosomal membrane and luminal proteins also tended to be higher in *TRIM46* knockout HeLa cells (Fig. 2C, D) and HEK293T cells (Fig. S2A, B). Thus, lysosomal abundance is increased in *TRIM46* knockout cells. To further assess lysosomal function in *TRIM46* knockout cells, we used DQ-BSA, a molecule that becomes fluorescent upon hydrolysis by lysosomal proteases in an acidic environment [34]. *TRIM46* knockout cells exhibited increased DQ-BSA signal relative to WT cells (Fig. S2C, D). However, when DQ-BSA signal was normalized to the abundance of lysosomes based on LAMP2 staining, *TRIM46* knockout cells were indistinguishable from WT (Fig. S2E). This indicates that the degradative functions of lysosomes are intact in *TRIM46* knockout cells. These data suggest that the increased lysosome abundance is not due to the accumulation of dysfunctional lysosomes, but instead due to lysosomal biogenesis.

**Figure 2.**
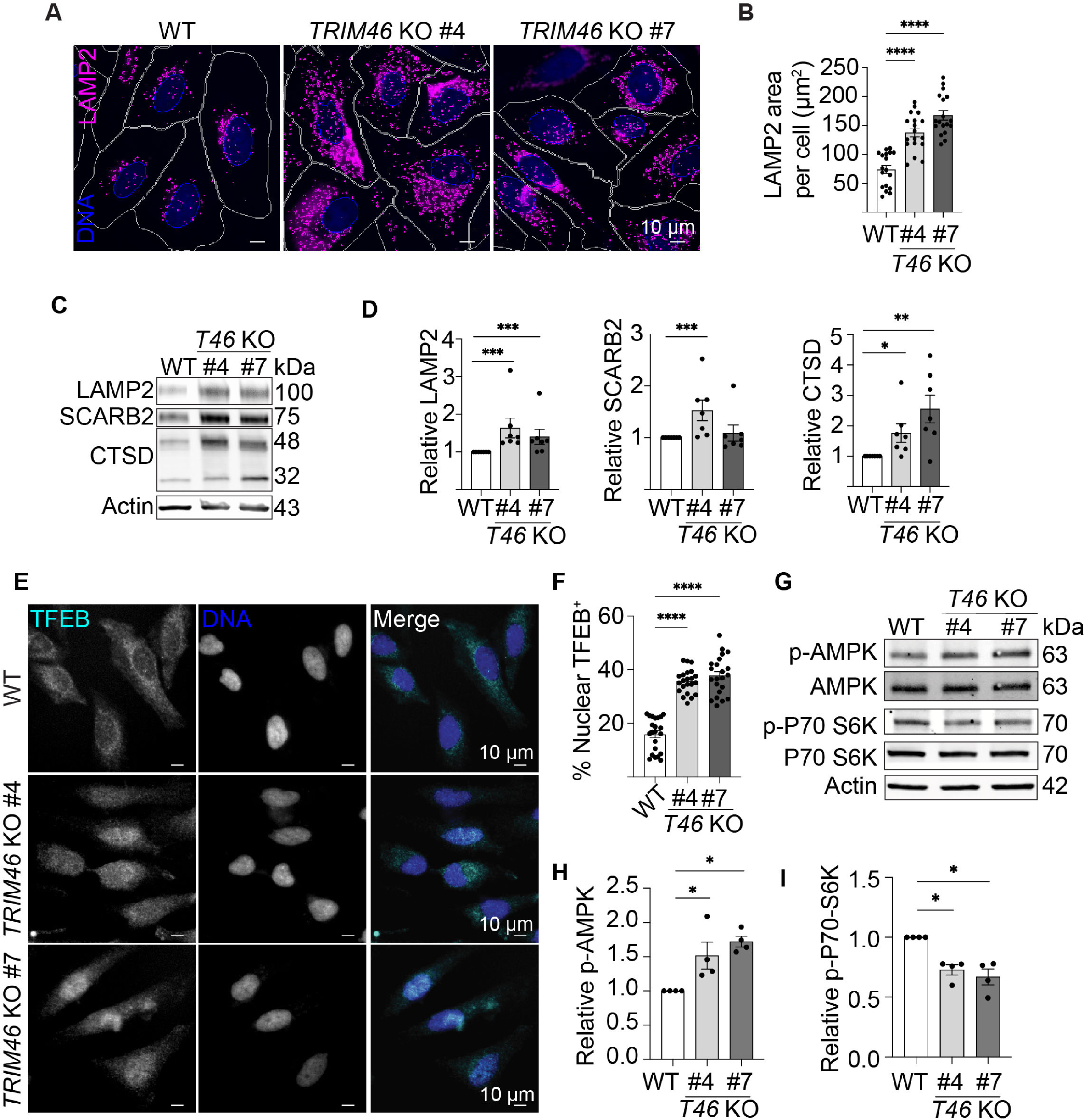
TRIM46 deficiency increases lysosome biogenesis. (**A, B**) High-content imaging analysis of LAMP2 abundance in WT HeLa and two *TRIM46* knockout clones. The average area of punctate LAMP2 per cell from >500 cells was quantitated and graphed in (B). (**C, D**) The levels of the indicated proteins in WT and *TRIM46* knockout cells were measured by immunoblotting (C) with quantitation from 6 independent experiments shown in (D). (**E, F**) High content image analysis of TFEB localization in WT and *TRIM46* knockout HeLa cells. The percentage of cells showing nuclear-localized TFEB was plotted in (F). Each data point represents the average calculated from >500 cells. (**G-I**) Immunoblot analysis of the effects of TRIM46 deficiency on MTORC1 and AMPK kinase activity. Quantification of relative protein abundance of phospho-AMPKα (T172) and phospho-P70-S6K (T389) are shown in (H) and (I), respectively. Each data point represents an independent experiment. Statistical analyses were performed using non-parametric Mann-Whitney U test (D, H, I) or ordinary one-way ANOVA followed by Dunnett’s multiple-comparison test (B, F). Data: mean ± SEM; *, p < 0.05; **, p < 0.01; ***, p < 0.001; ****, p < 0.0001.

Lysosomal biogenesis is controlled at the transcriptional level by the MiTF/TFE family of transcription factors, which includes the protein TFEB. TFEB shuttles between the cytoplasm and nucleus depending on its phosphorylation status to activate the expression of a large number of autophagy- or lysosome-related genes [35–37]. TFEB staining revealed that *TRIM46* knockout cells exhibited a nearly two-fold increase in TFEB nuclear localization compared to WT cells (Figure 2E, F), consistent with enhanced lysosomal biogenesis in *TRIM46* knockout cells. TFEB is under the control of the master metabolic regulating kinases MTORC1 and AMPK [35,38–41]. *TRIM46* knockout reproducibly increased the level of active phosphorylated AMPK while the phosphorylation of RPS6KB/p70 S6 kinase, a substrate of MTORC1, was decreased (Fig. 2G-I). Active MTORC1 is primarily localized on lysosomes [42]. In confocal microscopy experiments, we found that the lysosomal localization of MTOR was reduced in *TRIM46* knockout cells (Fig. S2F, G), further indicating reduced MTORC1 activation. Overall, these data show that TRIM46 restrains TFEB-mediated lysosomal biogenesis associated with MTORC1 inactivation.

### The Golgi apparatus is disrupted and subject to non-degradative Atg8ylation in *TRIM46 knockout cells*

We next considered how *TRIM46* deficiency triggered TFEB activation. As the master regulator of lysosome biogenesis, TFEB activation is strongly associated with lysosomal damage. However, our data shown in Fig. S2 indicates that the lysosomes in *TRIM46* knockout cells are functional. We thus considered whether damage to other organelles could explain the phenotypes observed in *TRIM46* knockout cells. Structural disruption of the Golgi has been reported to inactivate MTOR signaling while inducing Atg8ylation [9,43,44]. The architecture of the Golgi apparatus is maintained by microtubules [45,46]. TRIM46 localizes to microtubules and has been implicated in bundling axonal microtubules in neurons [27]. We thus hypothesized that Golgi structure might be disrupted in *TRIM46* knockout cells via microtubule disorganization, and that this may explain the MTOR and TFEB phenotypes that we have observed. In agreement with this hypothesis, we found that mCherry-tagged TRIM46 exclusively localized to microtubules when expressed in HeLa cells (Fig. S3A). The microtubules in *TRIM46* knockout cells were shorter and more disorganized than in WT cells (Fig. S3B-D). Along with the microtubule phenotypes, we observed that the Golgi apparatus in *TRIM46* knockout cells was enlarged and less coalesced, exhibiting a mixture of dispersed Golgi stacks and increased small puncta positive for TGOLN2 (trans-Golgi network protein 2; Fig. 3A). As a measure of Golgi fragmentation, we used high content imaging and analysis to selectively identify “small” TGOLN2-positive structures (smaller than 15 μm^2^) and quantitate them in WT and *TRIM46* knockout cells. Using this analysis, we saw that the number of Golgi fragments was increased by ∼2-fold in both *TRIM46* knockout cell lines relative to WT (Fig. 3B and S3E). Additionally, the total area of TGOLN2, including both small and large TGOLN2-positive structures, is increased in *TRIM46* knockout cells (Fig. 3C). Although imaging analysis revealed an increased Golgi area in *TRIM46* knockout cells, the abundance of Golgi-resident proteins including ACBD3/GCP60 and the *trans*-Golgi protein TGOLN2 was unchanged, and only a slight increase was observed in the endosomal/trans-Golgi network (TGN) localized protein WIPI1 (Fig. S3F). These findings suggest that the increased Golgi area reflects structural dispersal rather than an upregulation of Golgi protein synthesis in *TRIM46* knockout cells. Together, these data demonstrate that TRIM46 is essential for maintaining Golgi architecture, likely through TRIM46’s ability to organize microtubules.

**Figure 3.**
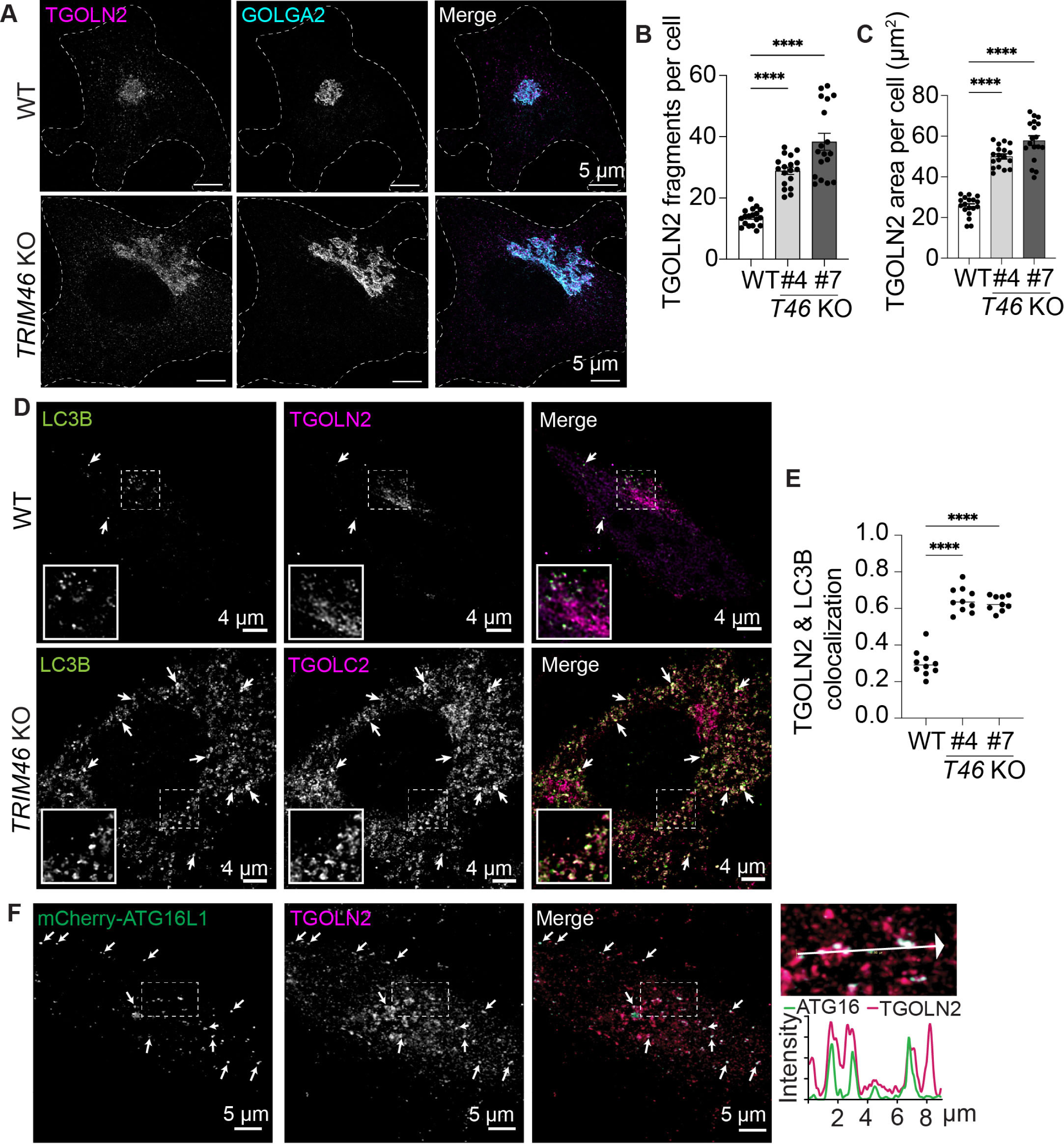
TRIM46 deficiency induces Golgi fragmentation and non-degradative Atg8ylation. (**A**) Maximum image projection confocal images of WT and *TRIM46* knockout HeLa cells stained with trans-Golgi marker TGOLN2 and cis-Golgi marker GOLGA2. (**B, C**) High-content imaging analysis of trans-Golgi network (TGN) fragmentation in WT and *TRIM46* knockout cells. Representative images are in Supplementary Figure S3E. TGN fragmentation was quantified by determining the number of “small” TGOLN2-positive structures (<15 μm²) per cell (B). The total area of TGOLN2-positive area was measured for individual cells, and the mean value of over 500 cells was plotted as a single data point (C). (**D**) Confocal analysis of colocalization between LC3B and TGOLN2 in WT and *TRIM46* knockout (shown, KO #4) HeLa cells. Zoomed in images of the boxed regions are shown below. Arrows indicate colocalized signal. (**E**) Colocalization between TGOLN2 and LC3B was quantified using Pearson’s correlation coefficient analysis. Each point represents data from a different confocal image. (**F**) Confocal analysis of mCherry-ATG16L1 and TGOLN2 localization in transiently transfected *TRIM46* knockout cells. Small arrows indicate colocalizing puncta. An enlarged image of the boxed region is shown. The large arrow indicates the path of the measured intensity profile. Each data point represents the average of >500 cells (B and C) or a single image field (E). Statistical analyses were performed using ordinary one-way ANOVA followed by Dunnett’s multiple-comparison test. Data: mean ± SEM; **, p < 0.01, ****; p < 0.0001.

In our siRNA screen (Fig. 1), we observed that *TRIM46* knockdown increased the abundance of non-acidified LC3B structures, suggesting that TRIM46 may inhibit a non-degradative Atg8ylation pathway. Several other groups have observed non-degradative conjugation of LC3 to Golgi membranes [9,44,47], and so we wondered if some of the LC3-positive structures seen in *TRIM46* knockout cells colocalized with the *trans*-Golgi marker TGOLN2. Indeed, we found substantial overlap between the small TGOLN2-positive structures and the Atg8 proteins LC3B and GABARAP in *TRIM46* knockout cells (Fig. 3D, E; S4A-E). Analysis of three-dimensional reconstructions of deconvolved confocal images of *TRIM46* knockout cells that had been probed with antibodies recognizing TGOLN2 and LC3B revealed that the LC3B-positive structures that colocalized with TGOLN2 had irregular morphologies that corresponded with the shape of the TGOLN2 structure (Fig. S4F). The shape of these LC3B-positive structures was different from what might be expected of autophagosomes, which tend to be round. In agreement with this observation, quantitative analysis showed that the LC3B-positive structures in TRIM46 knockout cells showed reduced sphericity when compared to LC3B-positive structures in WT cells (Fig. S4G). Subcellular fractionation experiments also indicated association between lipidated LC3B and TGOLN2-positive membranes (Fig. S4H). To enrich lysosomal and TGN vesicles, we performed sequential, differential centrifugation including ultracentrifugation at 100,000 g. The resulting pellets were lysed, and equal amounts of protein were analyzed. Fractionation was confirmed by immunoblotting with LAMP2 and TGOLN2 as lysosomal and TGN markers, respectively. In WT cells, lipidated LC3B was predominantly found in 21 k fraction, with substantially less LC3B-II cofractionating with TGOLN2 in the 100 k fraction. In contrast, roughly equal amounts of LC3B-II were found in the 21 k and 100 k fractions in *TRIM46* knockout cells (Fig. S4H). The findings presented above are consistent with direct conjugation of LC3B to the TGOLN2-positive membranes. In agreement with this concept, ATG16L1, a key component of the Atg8ylation machinery, shows strongly enhanced colocalization with TGOLN2 in *TRIM46* knockout cells (Fig. 3F and S4I-K). We next used mKeima-fused YIPF3 as a reporter to determine if the atg8ylated Golgi fragments were delivered to the lysosome for degradation in a selective autophagy-based process called ‘Golgiphagy’ [7,48,49]. YIPF3 is a Golgi-resident transmembrane protein that serves as a Golgiphagy receptor and is degraded by autophagy following amino acid starvation [48,49]. We saw no difference in the delivery of the mKeima-YIPF3 reporter to acidified compartments when we compared WT with *TRIM46* knockout cells, despite positive controls (amino acid starvation) and negative controls (BafA1) behaving as expected in this assay (Fig. S4L). These results, along with data showing that *TRIM46* knockout does not reduce the abundance of several Golgi-resident proteins (Fig. S3F), show that *TRIM46* knockout does not increase Golgiphagy, and instead imply that the observed Golgi Atg8ylation is non-degradative.

### *TRIM46* knockout activates CASM of Golgi membranes

Our data indicate that *TRIM46* knockout induces non-autophagy Atg8ylation of TGOLN2-positive membranes in a manner reminiscent of CASM. To further validate this, we inhibited or knocked down proteins that are essential for autophagy but are dispensable for CASM [50–52]. When WT cells are treated with VPS34-IN1, a compound that inhibits phagophore formation, LC3B-II levels were reduced by ∼60%. In contrast, this treatment only modestly impacted LC3B-II levels in *TRIM46* knockout cells (∼20% reduction; Fig. 4A, B). Next, we employed siRNA to knock down the expression of the core autophagy proteins ATG7, ATG13, BECN1, and ULK1 to determine how this impacted the elevated levels of LC3B-II and the increased abundance of LC3-positive structures in *TRIM46* knockout cells (Fig. 4C-F; S5A-F). As expected, knockdown of ATG7 reduced lipidated LC3B and LC3B puncta. ATG7 knockdown also reduced the colocalization between LC3B and TGOLN2 (Fig. 4G; S5C). However, knocking down the expression of ATG13 and BECN1 did not reduce the abundance of LC3B-II or LC3 puncta in *TRIM46* knockout cells. ULK1 knockdown seemed to reduce Atg8ylation in *TRIM46* knockout cells, but this trend was not statistically significant (Fig. 4E-G). These data confirm that much of the excessive Atg8ylation seen in *TRIM46* knockout cells is attributable to CASM and not autophagy.

**Figure 4.**
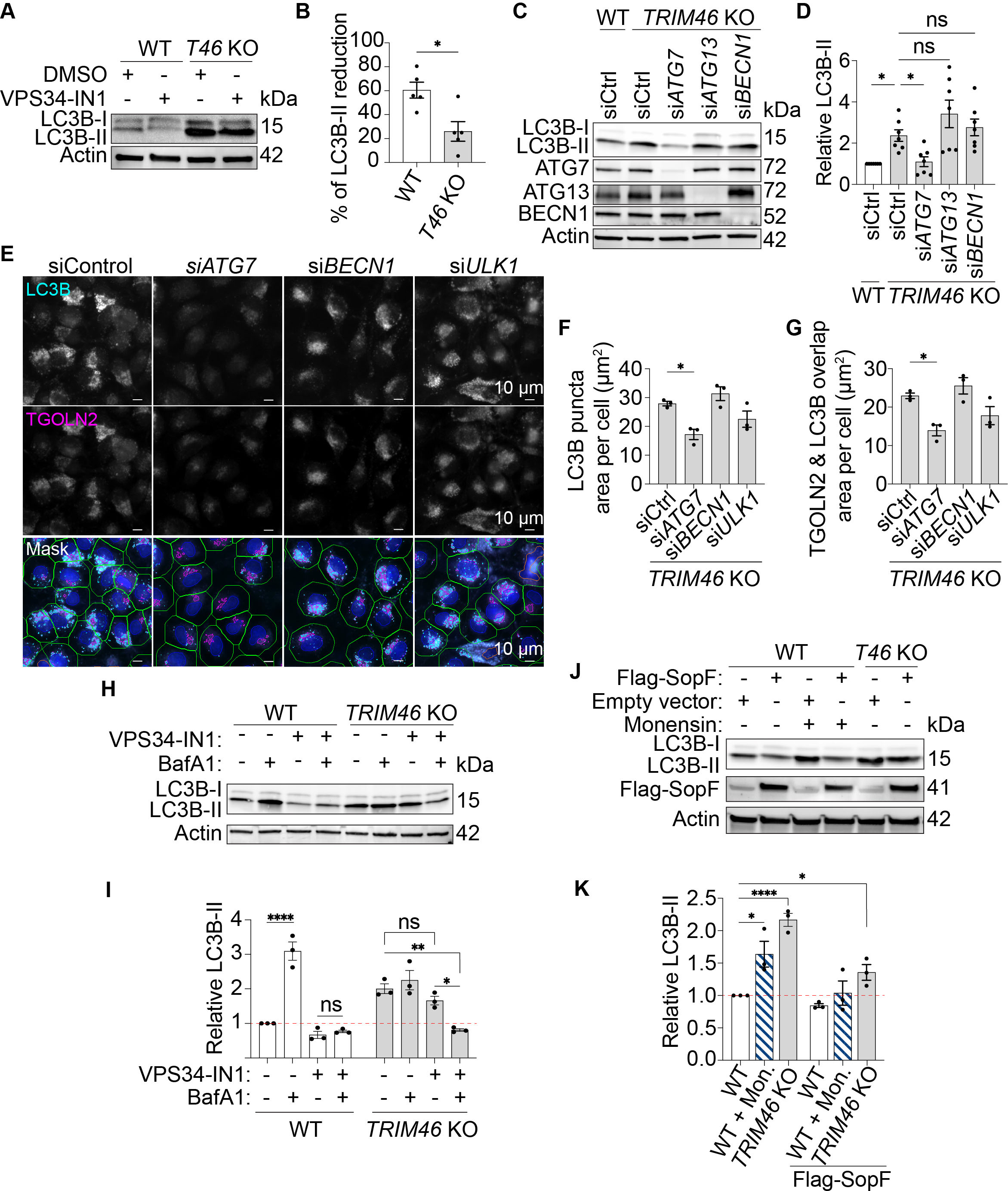
Enhanced Atg8ylation in *TRIM46* knockout cells is independent of upstream autophagy regulatory factors. (**A-B**) WT or *TRIM46* knockout HeLa cells were treated with 10 μM of VPS34-IN1 for 4 h prior to lysis and immunoblotting (A). Graph (B) shows the relative reduction, expressed as a percent, in LC3-II abundance following VPS34-IN1 treatment. Data points represent independent experiments. (**C-D**) WT or *TRIM46* knockout HeLa cells were transfected with the indicated siRNA prior to lysis, immunoblotting (C), and quantitation of LC3B-II levels relative to that seen in WT cells (D). Data points represent independent experiments. (**E-G**) Representative high content images of *TRIM46* knockout HeLa cells transfected with the indicated siRNA and stained with antibodies recognizing LC3B and TGOLN2. Plots show the abundance of punctate LC3B area per cells (F) and the overlapping area between TGOLN2 and LC3B (G). (**H**) Immunoblot analysis of the effect of VPS34-IN1 and BafA1 on LC3B lipidation in WT and *TRIM46* knockout cells. Cells were treated with inhibitors (VPS34-IN1: 10 μM, BafA1: 100 nM) or DMSO vehicle control for 4h. (**I**) Plot shows the abundance of LC3B-II relative to that seen in DMSO-treated WT cells. (**J**) Immunoblot analysis of the effect of SopF on LC3B lipidation in WT and *TRIM46* knockout cells. Cells were transfected with either an empty vector or Flag-SopF. 24 h post-transfection, cells were treated with either ethanol or 10 μM of monensin for 1 h. (**K**) Plot shows the abundance of LC3B-II relative to that seen in ethanol treated WT cells. Data points represent biological replicates, each based on an average of more than 500 cells for microscopy experiments. Statistical analyses were performed using an unpaired t-test (B), Kruskal-Wallis followed by Dunn’s multiple-comparison test (D), ordinary one-way ANOVA followed by Dunnett’s multiple-comparison test (F, G), or two-way ANOVA followed by Tukey’s multiple-comparison test (I, K). Data: mean ± SEM; *, p < 0.05; **, p < 0.01; ****, p < 0.0001; ns, not significant.

Lysosomal deacidification promotes a sub-type of CASM called V-ATPase-ATG16L1-induced LC3 lipidation (VAIL) [53,54]. In this pathway, assembly and activation of V-ATPase complexes on deacidified membranes recruit the ATG16L1 E3 ligase complex via direct protein–protein interaction [55,56]. Analogous to lysosomes, the lumen of the Golgi is more acidic than the surrounding cytosol, and so we asked whether the elevated Atg8ylation seen in *TRIM46*-knockout cells was dependent on V-ATPase assembly and/or VAIL. Pharmacological inhibition of V-ATPase assembly with BafA1 substantially reduced LC3B-II levels in *TRIM46* knockout cells cotreated with VPS34-IN1(Fig. 4H, I). We then tested whether expression of the bacterial effector SopF, which disrupts VAIL by blocking V-ATPase-ATG16L1 interactions [53,56], reduced LC3B lipidation in *TRIM46* knockout cells. As expected, SopF expression completely abrogated LC3B conversion induced by the VAIL-inducer monensin. In contrast, *TRIM46* knockout cells expressing SopF still showed substantially elevated levels of LC3B-II, albeit they were reduced relative to *TRIM46* knockout cells transfected with a control plasmid (Fig. 4J, K). Together, these findings indicate that Golgi membrane Atg8ylation in TRIM46-deficient cells requires V-ATPase activity but proceeds through a mechanism distinct from canonical VAIL.

### Impacts of CASM on TFEB activation and Golgi morphology in TRIM46-deficient cells

In most cases, the physiological consequences of CASM are not entirely clear. To investigate whether CASM contributes to the phenotypes observed in *TRIM46* knockout cells, we knocked down the expression of key components of the Atg8ylation machinery. siRNA-mediated knockdown of *ATG7* expression substantially reduced the percentage of *TRIM46* knockout cells that were positive for TFEB nuclear localization (Fig. 5A), implicating CASM in the increased lysosomal biogenesis seen in *TRIM46* knockout cells.

**Figure 5.**
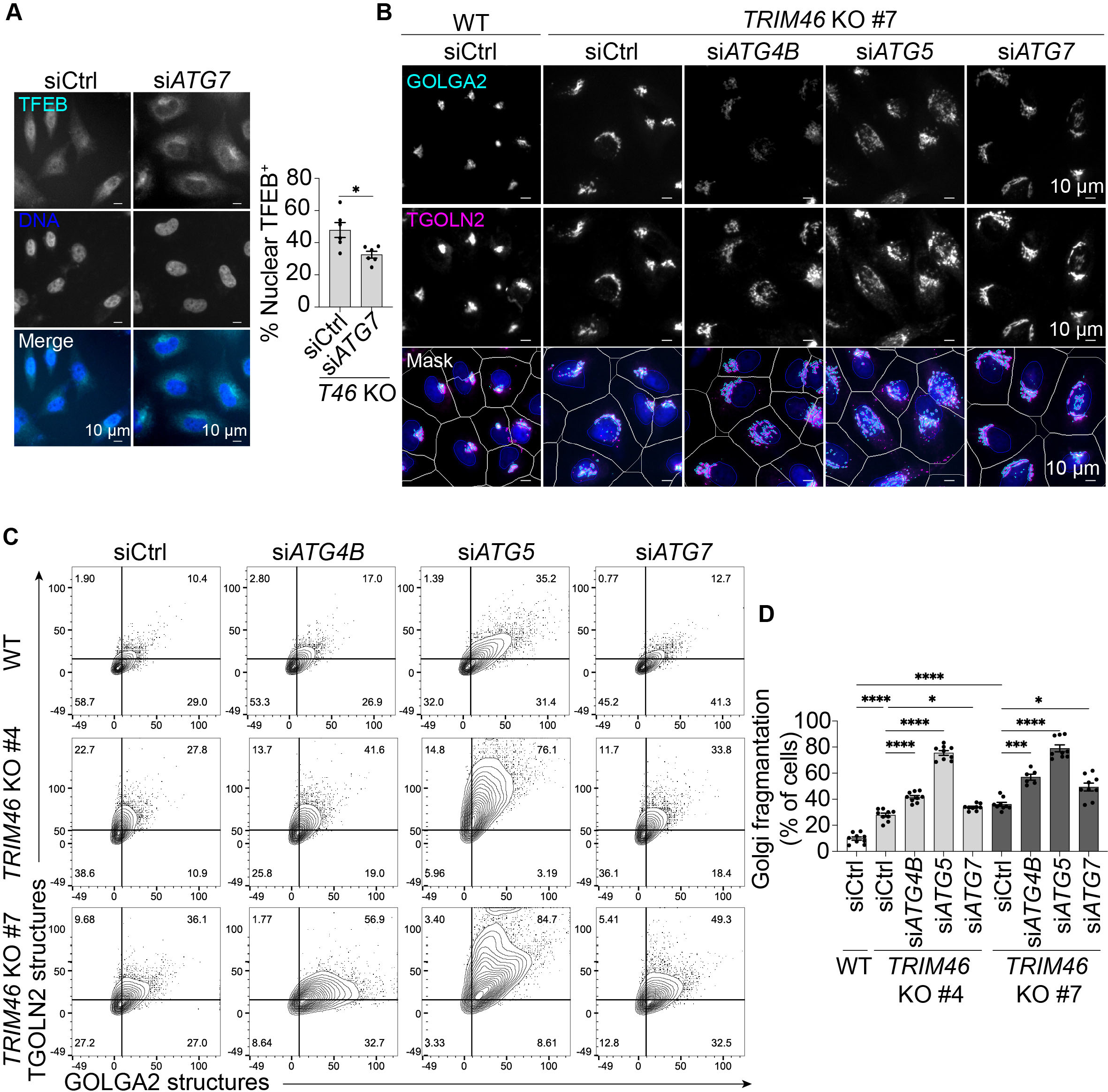
Atg8ylation machinery promotes TFEB activation and preserves Golgi architecture. (**A**) Representative images showing TFEB staining in *TRIM46* knockout HeLa cells transfected with control or *ATG7* siRNA (left). Quantification (right) shows the percentage of cells with nuclear TFEB localization as determined by high content imaging. Data points represent biological replicates; each calculated from the average of >500 cells. (**B**) Representative images of *TRIM46* knockout cells transfected with the indicated siRNA and stained with antibodies recognizing GOLGA2 and TGOLN2. (**C**) Contour plots show the number of distinct GOLGA2 and TGOLN2 structures per cell. The numbers in the upper right-hand quadrants indicate the percentage of cells exhibiting Golgi fragmentation, defined as >10 GOLGA2^+^ structures and >15 TGOLN2^+^ structures per cell. N ≥6030 cells analyzed per sample. (**D**) Percentage of cells classified as having fragmented Golgi. Data points represent biological replicates; each calculated from the average of >500 cells. Statistical analyses were performed using an unpaired t-test (A), Welch’s ANOVA followed by Dunnett’s multiple-comparison test (D). Data: mean ± SEM; *, p < 0.05; ***, p < 0.001; ****, p < 0.0001.

Our data also demonstrate a role for CASM in regulating Golgi architecture in *TRIM46* knockout cells. As shown in Figure 3, the distribution of Golgi markers GOLGA2/GM130 and TGOLN2 was altered in *TRIM46* knockout cells. To determine the impact of Atg8ylation on Golgi architecture, we used high content microscopy to quantify the abundance of GOLGA2^+^ and TGOLN2^+^ structures in cells transfected with control siRNA or siRNA targeting *ATG4B*, *ATG5*, and *ATG7*; factors that are required for all Atg8ylation processes including CASM. Relative to WT, *TRIM46* knockout cells exhibited increased abundance of GOLGA2^+^ and TGOLN2^+^ structures when transfected with non-targeting siRNA, indicative of increased Golgi fragmentation. Remarkably, the Golgi fragmentation seen in *TRIM46* knockout cells was substantially exacerbated when the Atg8ylation machinery was inhibited, with *TRIM46* knockout cells transfected with siRNA targeting the three ATGs having a substantially increased number and cross-sectional area of GOLGA2^+^ and TGOLN2^+^ structures (Fig. 5B-D and S6A, B). Morphologically, *ATG4B*, *ATG5*, and *ATG7* knockdown in *TRIM46* knockout cells all promoted the loss of Golgi ribbon structures and correspondingly increased numbers of discrete dispersed ministacks (Fig. 5B). Collectively, these data show that Atg8ylation contributes to repairing or maintaining Golgi architecture under stress conditions. Interestingly, we observed that *ATG5* knockdown also significantly increased Golgi fragmentation in WT cells (Fig. 5C). Knockdown of *ATG4B* and *ATG7* also followed this trend, but the effects were not statistically significant. These data suggest that Atg8ylation may contribute to Golgi architecture under normal growth conditions. The ability of Atg8ylation to maintain Golgi architecture is likely mediated by CASM, since autophagy inhibition with VPS34-IN1 did not exacerbate the Golgi fragmentation phenotype in *TRIM46* knockout cells, and in fact had the opposite effect (Fig. S6C-E). SopF expression had no impact on Golgi morphology in *TRIM46* knockout cells (Fig. S6F). Taken together, these results demonstrate a novel role of CASM in supporting Golgi ribbon formation and limiting Golgi structural disruption under stress.

### Chemical disruption of the Golgi architecture phenocopies *TRIM46* knockout

We have demonstrated that TRIM46 deficiency leads to disorganization of the microtubule network, Golgi fragmentation, and CASM activation. To determine whether microtubule disorganization alone is sufficient to mimic the phenotypes seen in *TRIM46* knockout cells, we treated WT HeLa cells with the microtubule-depolymerizing agents nocodazole and vinblastine. As expected, both treatments significantly increase TGOLN2-positive fragments (Fig. 6A, B and S7A, B). Similar to *TRIM46* knockout cells, both nocodazole- and vinblastine-treated cells showed a significant increase in LC3B puncta and LC3B/TGOLN2 colocalization, suggesting CASM activation (Fig. 6C-E; S7C-E). Microtubule disruption also increased lysosome abundance (Fig. 6F, G; S7F, G) and increased TFEB nuclear localization (Fig 6H, I; S7H, I), mirroring the effects observed in *TRIM46* knockout cells. Thus, disorganization of the microtubule network, whether by drug treatment or by TRIM46 deficiency, results in CASM and lysosomal biogenesis.

**Figure 6.**
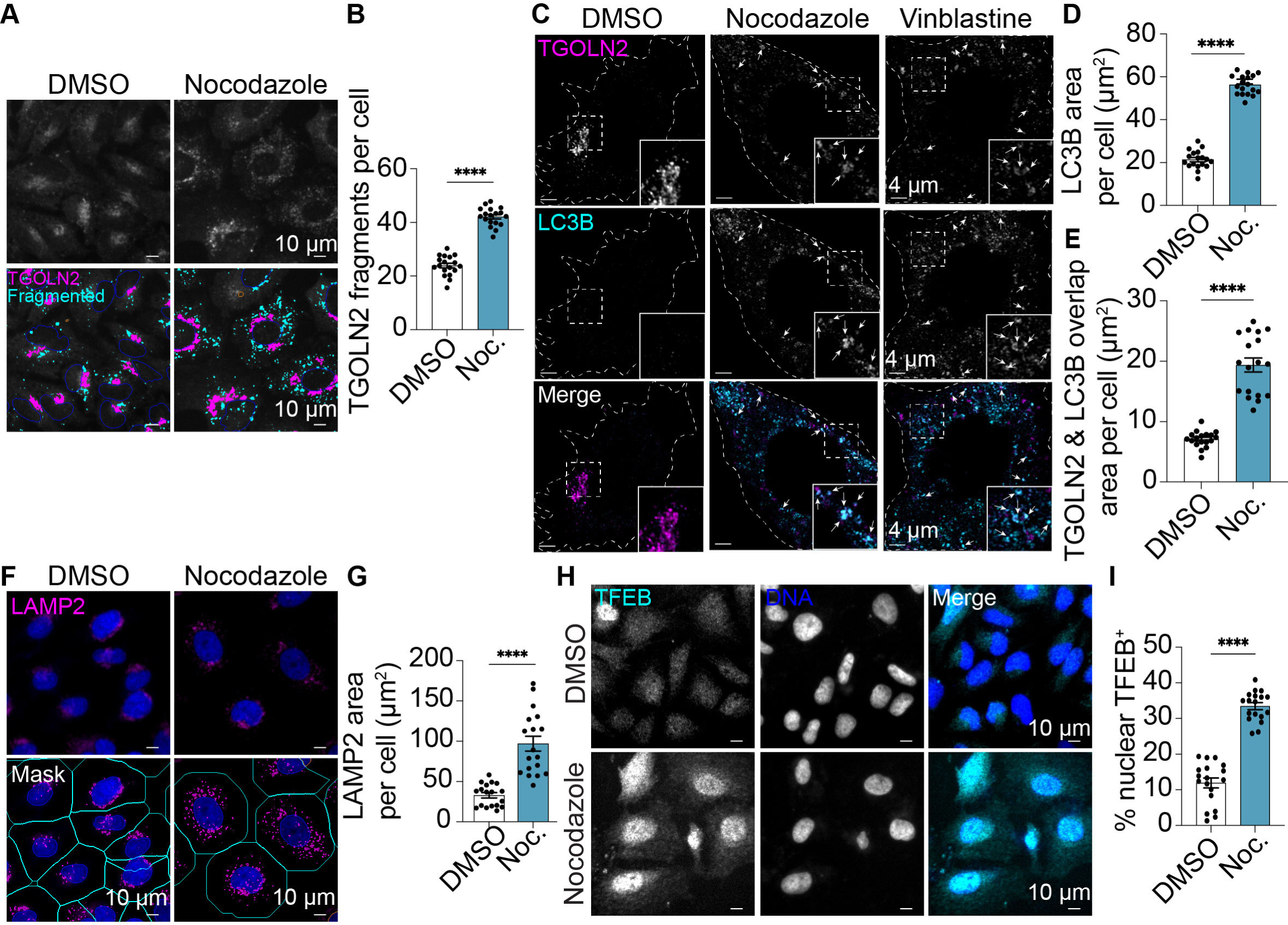
Microtubule disruption phenocopies TRIM46 deficiency, triggering Golgi Atg8ylation and increased lysosomal biogenesis. WT HeLa cells were treated with 100 nM of nocodazole for 16 h. (**A-B**) High-content imaging analysis of TGN fragmentation measured by counting TGOLN2-positive fragments smaller than 15 μm² (cyan mask). Magenta mask indicates TGOLN2-positive structures with surface areas >15 μm². (**C**) Confocal microscopic analysis of TGOLN2 and LC3B colocalization in HeLa cells treated with nocodazole, vinblastine, or DMSO control. Arrows indicate colocalizes structures. Insets show magnified images of the area within the dashed-line boxes. (**D-E**) Plots show coalesced LC3-positive area per cell (D), and overlapping area between TGOLN2 and LC3B (E), quantified by high-content microscopy imaging analysis following nocodazole treatment. (**F-G**) High-content imaging analysis of WT HeLa cells of LAMP2 abundance following nocodazole treatment. (**H-I**) High content imaging analysis of TFEB nuclear localization following nocodazole treatment. Statistical analyses were performed using an unpaired t-test. Data: mean ± SEM; ****, p < 0.0001. Each data point represents the average of more than 500 cells.

We next asked whether we could recapitulate the phenotypes seen in *TRIM46* knockout cells by disrupting Golgi apparatus architecture without targeting microtubules. To test this, we treated HeLa cells with brefeldin A, a compound that induces Golgi disassembly by interfering with the formation of COP-I vesicles. As expected, brefeldin A increased the number of Golgi fragments per cell in WT HeLa cells (Fig. S7J, K). Additionally, we found that it significantly increased the abundance of LC3B puncta per cell and the colocalization between LC3B and TGOLN2 (Fig. S7L-N). Brefeldin A treatment also resulted in a >2-fold increase in TFEB nuclear localization (Fig. S7O, P). These data show that Golgi Atg8ylation and TFEB activation are generalized cellular responses to disruption of Golgi architecture.

### Knockdown of Atg8ylation factors impedes the repair of acutely fragmented Golgi structures

Data shown in Figure 5 demonstrated that CASM supports Golgi architecture in the setting of chronic microtubule disorganization caused by TRIM46 deficiency. However, those experiments could not distinguish whether CASM maintains Golgi organization or whether it actively promotes Golgi repair following fragmentation. To directly test a role in repair, we employed an acute fragmentation model in which Golgi fragmentation is induced by nocodazole and reassembly is monitored following drug washout (Fig. 7A, B). Golgi morphology was quantified by high content imaging in HeLa cells transfected with control siRNA or siRNA targeting Atg8ylation factors *ATG4B*, *ATG5*, and *ATG7* prior to and during Golgi fragmentation with nocodazole or at different time points following nocodazole washout. In our assay, nocodazole treatment caused Golgi fragmentation in ∼90% of cells. In control cells, Golgi reassembly was evident as soon as 15 min after nocodazole washout and was nearly complete by 2 h. In contrast, the restoration of normal Golgi architecture was impaired in cells subjected to knockdown of any Atg8ylation factor, with significantly more cells showing a fragmented Golgi phenotype at 15, 30, and 60 min after washout. *ATG5* knockdown produced the strongest and most durable defect, with an elevated number of cells showing Golgi fragmentation up to 2 h after nocodazole removal (Fig. 7A, B). Together, these findings demonstrate that CASM is required for efficient repair of acutely fragmented Golgi structures and provide a framework for interpreting the *TRIM46* knockout phenotype: ongoing CASM-mediated Atg8ylation may continuously attempt to restore Golgi organization under chronic microtubule disruption, such that disrupting Atg8ylation further exacerbates Golgi fragmentation.

**Figure 7.**
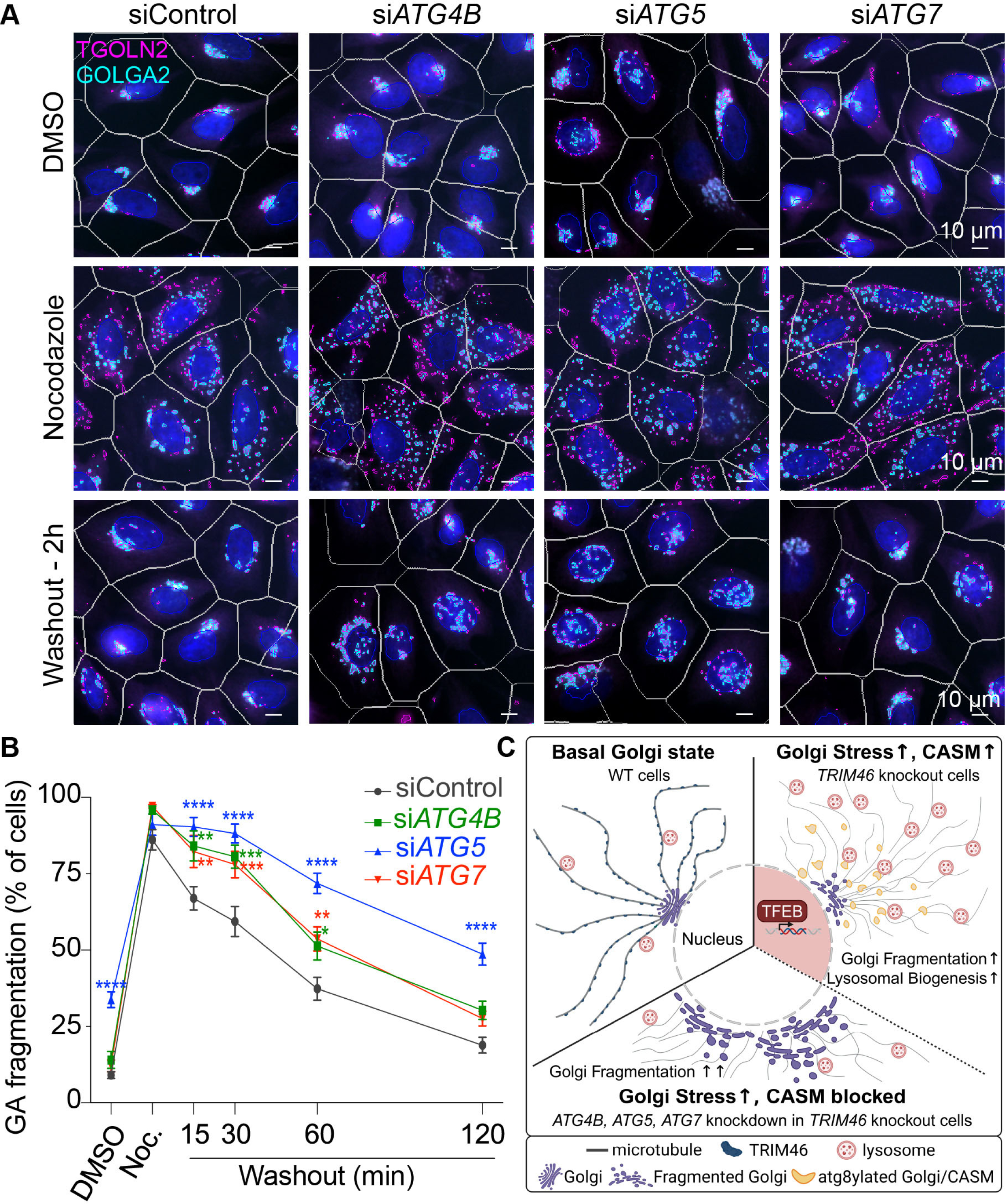
Knockdown of Atg8ylation factors delays Golgi repair. WT HeLa cells were transfected with control siRNA or siRNAs targeting *ATG4B*, *ATG5*, or *ATG7*. 72 h post-transfection, cells were treated with 500 nM nocodazole for 4 h, followed by washout and incubation in nocodazole-free media for 15, 30, 60 or 120 min. Cells were then fixed and stained for GOLGA2 and TGOLN2, and the extent of Golgi reassembly was quantified by counting cells with >10 GOLGA2^+^ structures and >15 TGOLN2^+^ structures per cell. A total of ≥14517 cells per sample were analyzed from 4 independent experiments. (**A**) Representative images of *TRIM46* knockout cells transfected with the indicated siRNAs, treated with nocodazole, and subjected to a 2-h washout. Cells were stained with antibodies against GOLGA2 and TGOLN2. (**B**) Line graphs show the rate of Golgi reassembly at different time points after nocodazole washout in the indicated knockdown conditions. (**C**) Role of CASM in restoring Golgi architecture in the absence of TRIM46. Top left panel, TRIM46 localizes to microtubules and maintains the organization of microtubule fibers. Top right panel, in the absence of TRIM46, microtubules are disorganized, leading to Golgi fragmentation. In response, v-ATPase subunits are assembled on Golgi-membranes, triggering CASM and consequent TFEB activation and lysosomal biogenesis. Bottom panel, CASM inhibition by knockdown of the Atg8ylation machinery in *TRIM46* knockout cells results in increased Golgi fragmentation and attenuated TFEB activation. Statistical analyses were performed using two-way ANOVA followed by Tukey’s multiple-comparison test. Data: mean ± SEM; *, p < 0.05; **, p < 0.01; ***, p < 0.001; ****, p < 0.0001.

## Discussion

Golgi architecture disruption is associated with multiple pathological conditions including viral infection, cancer, and neurodegenerative disease [57–61]. How cells respond to or repair these architectural changes remains unclear. Our study has shown that disruption of the Golgi apparatus activates a generalized response including Golgi Atg8ylation and TFEB-driven lysosomal biogenesis, with CASM enabling the restoration of Golgi architecture following acute or chronic fragmentation (Fig. 7C). We started our studies focusing on TRIM46, which likely alters Golgi structure indirectly through its actions in establishing microtubule organization. Supporting this concept, we found that treatment of wild type cells with the microtubule depolarizing compounds vinblastine or nocodazole phenocopied what we observed with the *TRIM46* knockout cells. Interestingly, brefeldin A, which disrupts Golgi structure and function without impacting microtubule organization, also induced Golgi Atg8ylation and TFEB activation. This result suggests that loss of Golgi architecture, rather than disruption of microtubule organization, is the primary trigger for these phenotypes. In agreement with this concept, blockage of post-Golgi trafficking by overexpression of the secreted protein DLK1[9], downregulation of a Golgi-localized membrane tether [62], or various Golgi-damaging treatments [44,63] all induce LC3 conversion.

Key questions remain about why cells engage the Atg8ylation machinery in response to Golgi or other membrane stress. We found that TRIM46 deficiency constitutively activated non-degradative Golgi Atg8ylation (CASM), and that CASM played an important role in mitigating the Golgi fragmentation seen in TRIM46-deficient or nocodazole-treated cells. Interestingly, we saw hints that the Atg8ylation machinery in general, and ATG5 in particular, may also contribute to Golgi architecture under normal homeostatic conditions. At the same time, our data disfavor models in which CASM enables increased Golgi biosynthesis or the delivery of new membrane to damaged Golgi structures, since these mechanisms would be expected to increase the size of the Golgi apparatus in an Atg8ylation-dependent manner, which is the opposite of what we found. While further mechanistic investigations are needed, our study expands the functional scope of CASM beyond previously described membrane perforation settings and identifies it as an active contributor to restoring complex of Golgi organization.

We also found that the increased activation of the transcription factor TFEB seen in *TRIM46* knockout cells was reversed following CASM inhibition. This result is consistent with several other recent findings that collectively reveal a positive feedback loop between Atg8ylation and TFEB activation [10,11]. Golgi damage, by either TRIM46 deficiency or by chemical disruption of the Golgi network, strongly promotes TFEB activation to enhance lysosomal biogenesis and upregulate the expression of proteins involved in CASM and autophagy. We speculate that the TFEB-dependent programs triggered by Golgi damage may function to counter the source of Golgi fragmentation or possibly to prepare the cell to initiate Golgiphagy should the extent of Golgi damage become more severe. This model conceptually aligns with what is known about lysosomal membrane permeabilization, in which CASM is activated in response to mild lysosomal damage while lysophagy, the autophagic targeting of damaged lysosomes, is reserved for lysosomes that are extensively damaged [64]. It is possible that a similar relationship exists with Golgi membranes: while *TRIM46* knockout does not cause sufficient damage to induce Golgiphagy, complete disruption of Golgi architecture with Brefeldin A is reported to induce autophagic degradation of Golgi-localized proteins [63].

Our studies, together with other reports of Golgi Atg8ylation in response to a variety of Golgi damaging conditions, raise the question of how the cell senses Golgi damage to activate Atg8ylation. However, whether distinct forms of Golgi damage converge on a common sensing pathway or instead trigger different danger signals remains unknown. Since the Golgi is a slightly acidic organelle, it is possible that some of the aforementioned Golgi damaging conditions could lead to proton leak and consequent deacidification. In endolysosomal membranes, this will lead to a V-ATPase-dependent subtype of CASM called VAIL, in which active V-ATPase assemblies recruit ATG16L1-containing E3 ligase complexes [53]. VAIL has been reported to occur on Golgi membranes in cells treated with the compound AMDE-1 [65]. Our data demonstrating that *TRIM46* knockout increases ATG16L1 recruitment to Golgi membranes and that V-ATPase inhibition with BafA1 reduces LC3 lipidation in *TRIM46* knockout cells are partially consistent with this model. However, experiments using SopF to directly test the role of V-ATPase-ATG16L1 complex formation in TRIM46 deficiency-driven Atg8ylation and in CASM-mediated Golgi repair yielded negative results. These findings indicate that Golgi fragmentation in *TRIM46* knockout cells activates CASM in a manner that appears to be dependent on V-ATPase activity but does not require the VAIL pathway. Thus, there is a need for future studies to determine how distinct Golgi fragmenting conditions recruit the Atg8ylation machinery and to uncover the mechanistic basis for how this machinery enables the reformation of Golgi architecture following stress.

Lysosomal deacidification also results in MTORC1 inactivation [42]. Interestingly, several studies have indicated that a pool of MTORC1 resides on the Golgi in addition to its well-established localization on lysosomes [62,66,67]. We found that MTORC1 activity was inhibited in *TRIM46* knockout cells, suggesting similar damage-responsive mechanisms between Golgi and lysosomes. Inhibition of MTORC1 activity can then result in or enhance autophagy and lysosomal biogenesis while also altering Golgi architecture [67,68].

Given the broad functions of TRIMs in autophagy, our initial goal was to screen TRIMs for roles in the late stages of autophagy. Follow-up studies revealed that TRIM46 deficiency activates a non-degradative form of Atg8ylation consistent with CASM. In addition to TRIM46, the knockdown of five other TRIMs also showed reduced LC3B acidification in our screen (Fig. 1). Further investigations are warranted to determine whether these TRIMs regulate Atg8ylation pathways indirectly, as in the case with TRIM46, or if they directly regulate autophagosome maturation.

In conclusion, our study identifies TRIM46 as a regulator of microtubule stability whose loss destabilizes the Golgi ribbon and triggers a coordinated membrane stress program characterized by CASM activation, lysosomal biogenesis, and altered MTORC1 and AMPK signaling. Our findings extend the growing body of literature on Golgi Atg8ylation by demonstrating that CASM is not merely associated with Golgi membranes, but actively contributes to the restoration of Golgi architecture following fragmentation. These findings raise the possibility that CASM-mediated Golgi repair contributes to cellular adaptation in neurodegenerative diseases and other pathological settings characterized by Golgi fragmentation.

## Materials and methods

See Table 1 for a list of key reagents used in this study.

### Antibodies

Primary antibodies were obtained as follows: mCherry (ab183628), GABARAP (ab109364; 1:500 for IF), and NPC2 (ab218192; 1:1,000 for WB) from Abcam; PRKAA/AMPK (2532; 1:1000 for WB), phospho-PRKAA/AMPK T172 (2535; 1:1,000 for WB), ATG13 (13468; 1:1,000 for WB), BECN1 (3459; 1:1,000 for WB), GOLGA2/GM130 (12480; 1:500 for IF, 1:1,000 for WB), MTOR (2983; 1:500 for IF), phospho-PRS6KB/p70 S6K (9205; 1:1,000 for WB), TFEB (4240; 1:500 for IF), and ULK1 (8054; 1:1,000 for WB) from Cell Signaling Technology; ACTB/actin (sc-58673; 1:1,000 for WB), Cas9 (sc-517386, 1:1000 for WB), LAMP2 (sc-18822; 1:1,000 for WB, 1:500 for IF), RPS6KB/p70 S6K (sc-8418; 1:1,000 for WB), and TUBA/alpha-tubulin (sc-23948; 1:500 for IF) from Santa Cruz Biotechnology; CTSD/cathepsin D (21327-1-AP, 1:1,000 for WB), MYC (16286-1-AP: 1:1,000 for WB), SCARB2/LIMP2 (27102-1-AP, 1:1,000 for WB), TRIM46 (21026-1-AP, 1:1,000 for WB) from Proteintech; LC3B (PM036; 1:500 for IF) from MBL and LC3B (L7543; 1:2,000 for WB) from Sigma; SQSTM1/p62 (610833; 1:2,000 for WB) from BD; ATG7 (MA5-32221; 1:1,000 for WB), ACBD3/GCP60 (MA5-25999; 1:1,000 for WB), TGOLN2/TGN46 (MA5-37930; 1:1,000 for WB), and WIPI1 (PA5-34973; 1:1000 for WB) from Thermo Fisher and TGOLN2 (AHP500G; 1:500 for IF) from Bio-Rad. Secondary antibodies were purchased as follows: anti-rabbit Alexa Fluor 488 (A11008; 1:1,000 for IF), anti-sheep Alexa Fluor 568 (A21099; 1:1,000 for IF), anti-rabbit Alexa Fluor 647 (A21244; 1:1,000 for IF), and anti-mouse Alexa Fluor 647 (A21235; 1:1,000 for IF) from Thermo Fisher; anti-mouse IRDye 680LT (925-68020; 1:10,000 for WB) and anti-rabbit IRDye 800CW (926-32211; 1:5,000 for WB) from LI-COR.

### Cell culture and treatment

All cell lines were obtained from ATCC and maintained in Dulbecco’s modified Eagle’s medium (DMEM; Thermo Fisher, 11965084) supplemented with 10% fetal bovine serum (FBS; Thermo Fisher, 26140095,) and 100 U/mL penicillin-streptomycin (Thermo Fisher, 15140122) in a humidified incubator at 37°C with 5% CO_2_. Bafilomycin A_1_ (Invivogen, tlrl-baf1; 100 μM), brefeldin A (Sigma, B6542; 5 mM), monensin (Sigma, M5273; 10μM), nocodazole (Sigma, M1404; 5 mM), Vps34-IN1 (Cayman chemical; 10 mM), vinblastine (Cayman chemical, 11762; 100 μM) were prepared as stock solution in DMSO at the indicated stock concentration. To measure autophagy flux and CASM in *TRIM46* knockout cells, cells were treated with 100 nM bafilomycin A1 with or without 10 μM vps34-IN1 for 4 or 6 h. For experiments examining TFEB nuclear localization or LC3B and LAMP2 induction, cells were treated with 100 nM brefeldin A, nocodazole, or vinblastine for 16 h.

### TRIM family siRNA screen

The methods employed for this experiment, and the results from analyzing eYFP signal on its own, were previously published [30]. Briefly, HeLa cells stably expressing mCherry-eYFP-LC3B [31] were plated into 96-well plates containing pre-plated siRNA smart pools and transfection reagent (Dharmacon). Cells were fixed 48 h after plating, stained with Hoechst 33342, and then imaged. Cell boundaries were determined based on nuclear staining with eYFP-positive and mCherry-positive LC3B puncta detected based on pre-set parameters in the iDev software. LC3B acidification was determined by subtracting the ratio of punctate eYFP-LC3B signal (neutral pH) to that of mCherry-LC3B signal (total LC3B) from 2; thus, values closer to 2 have increased delivery of the LC3B reporter to the lysosome while cells with inhibited delivery of LC3B to lysosomes will have reduced values. Those TRIMs whose LC3B acidification had values >3 standard deviations below the mean of cells transfected with non-targeting siRNA were considered hits. >500 cells per siRNA were analyzed in two independent experiments.

### Cloning

pDONR221-hTRIM46 (DNASU, HsCD00862242) was used as the donor for cloning human *TRIM46* into Gateway-compatible destination vectors, generating MYC- or mCherry-tagged *TRIM46* using LR Clonase (Thermo Fisher, 11791020). All Plasmid constructs were verified by whole-plasmid sequencing. For addback experiments, the PAM site in MYC- or mCherry-tagged *TRIM46* was disrupted to prevent degradation by constitutively expressing Cas9 in *TRIM46* knockout cells by introducing a silent mutation (1407G→T) using a site-directed mutagenesis kit (Agilent, 210518).

### Codon optimization and plasmid construction of SopF

The SopF coding sequence (Entrez Gene ID: STM1239) derived from *Salmonella enterica* was codon-optimized for human expression using the VectorBuilder codon optimization tool (VectorBuilder Inc.). The optimized sequence was synthesized and subcloned into the indicated expression vector (pRP[Exp]-EGFP). An N-terminal 3×FLAG tag was fused to SopF via a 3×GGGGS flexible linker. The plasmid contains two independent CMV promoter-driven expression cassettes encoding FLAG-SopF and EGFP separately; EGFP was used as a transfection control and is not fused to SopF. Co-expression of FLAG-SopF and EGFP from the dual CMV promoter construct was verified by anti-FLAG immunoblotting and fluorescence microscopy.

### Development of knockout or overexpressing cell lines

Knockout and stably overexpressing cell lines were generated by lentiviral transduction. Lentivirus was produced by co-transfecting HEK293T cells with pMD2.G, pPAX2, and pLV[CRISPR]-hCas9:T2A:Puro-U6-h*TRIM46* (VectorBuilder, VB900138-6978huh) or pLEX307-Halo-LC3 at a 2:3:5 ratio using the ProFection Mammalian Transfection System (Promega). After 48 h, virus-containing supernatants were collected, cleared of residual mammalian cells by centrifugation at 500 × g for 10 min, and filter through a 0.45 μm vacuum filter (SIGMA, SE1M003M00). Target cells (HeLa or HEK293T) were incubated with viral supernatants diluted in DMEM for 48 h and subsequently selected with 1 μg/mL puromycin (Thermo Fisher, J61278.MB). For *TRIM46* knockout cells, single-cell clones were isolated, and knockout was validated using the Guide-it Complete sgRNA Screening System (Takara, 632636). Briefly, sgRNA targeting *TRIM46* was generated using a customized forward PCR primer containing T7 promoter sequence, *TRIM46* target sequence, and Cas9 scaffold templates with the Guide-it sgRNA *In Vitro* Transcription Kit (Takara). The sgRNA template was validated on a 2% agarose gel, which showed a single band of ∼130 bp. sgRNA was synthesized by *In Vitro* transcription using the validated template. Genomic DNA was extracted from WT or *TRIM46* knockout cells using a genomic DNA extraction kit (QuantaBio, 95213-050), and the target region containing the CRISPR-Cas9 cleavage site was amplified by PCR. After purification of PCR fragments, the Cas9 cleavage assay was performed. The reaction products were loaded on a 1.5% agarose gel and imaged using a ChemiDoc Imaging System (Bio-Rad). For Halo-LC3B expressing HeLa wild type or *TRIM46* knockout cells, Halo-LC3B expression was confirmed by immunoblotting with anti-HaloTag antibody.

### RNA interference and transfection

Plasmid transfections were performed using Lipofectamine 2000 (ThermoFisher). Small interfering RNAs (siRNAs) were purchased from GE Dharmacon and were supplied as pools of four different siRNA oligos against the same mRNA. siRNA transfections were performed using Lipofectamine RNAiMAX (ThermoFisher) according to the manufacturer’s instructions.

### Nocodazole washout

WT HeLa cells were transfected with control siRNA or siRNAs targeting *ATG4B*, *ATG5*, or *ATG7* in 6-well plates. 48 h post-transfection, cells were re-seeded at a density of 1 × 10^4^ cells per well in 96-well plates. The following day, cells were treated with DMSO or 500 nM nocodazole for 4 h, followed by washout and incubation in nocodazole-free medium for 15, 30, 60, or 120 min. After incubation, cells were subjected to immunofluorescence staining as described below, and image analysis was performed using high-content microscopy.

### Immunofluorescent staining

For high content imaging, cells were seeded in either 24-well or 96-well plates (2 × 10⁴ cells per well for a 24-well plate or 1 × 10^4^ cells per well for a 96-well plate). For confocal experiments, 1 × 10^5^ cells were plated onto coverslips in 12-well plates. Cells were fixed with 4% paraformaldehyde for 3 min and then washed twice with 1 × PBS. In general, permeabilization was performed using 0.2% saponin and 1% BSA in PBS for 30 min, but for TFEB staining cells were permeabilized with 0.1% Triton X-100 and 0.1% Tween 20 for 10 min, followed by two washes with PBS and blocking with 1% BSA in PBS for 30 min. Primary antibodies were applied at the dilutions specified above and incubated with the cells for 1 h at room temperature. Following two washes with PBS to remove residual primary antibody solution, cells were incubated with appropriate secondary antibodies for 1 h at room temperature. Samples were then washed twice with 1 × PBS. Coverslips were mounted with ProLong Diamond Antifade mounting media (Invitrogen, P36970), while cells in multi-well plates were left in PBS for high content imaging.

### High content microscopy and analysis

Following immunofluorescence staining as described above, cells were counterstained with Hoechst 33342 to visualize nuclei. High-content imaging was performed using a Cellomics HCS or CellInsight CX7 instruments and analyzed with iDEV software (ThermoFisher). Hoechst 33342 nuclear staining was utilized for autofocusing, and regions of interest (ROIs) were defined relative to nuclear positions. Targets were identified based on fluorescence intensity within the ROI, and parameters such as object count, total area, total intensity, average intensity, co-localization area, and correlation coefficient between targets were analyzed on a per-cell basis using automated algorithms.

#### Puncta identification and quantitation, colocalization analysis, and analysis of nuclear localization

Puncta detection, colocalization, and nuclear localization were analyzed using the Cellomics colocalization BioApplication. Cell boundaries were defined based on either CellMask staining (Thermofisher, H32722) or the nuclei stained with Hoechst 33342 (Thermofisher, H3570). When nuclei were used to define cell boundaries, the nuclear parameter was extended to the surrounding cytoplasmic region. Single-cell segmentation was performed using nuclear intensity-based methods, and nuclei with irregular morphology or insufficient intensity were excluded to minimize segmentation errors. For puncta identification and quantification as well as colocalization analysis, the region of interest (ROI) was set to the cytoplasm, whereas for nuclear localization analysis, only the nuclear ROI was used. Target signals were measured within the defined ROI. Colocalization was quantified by automated calculation of the overlapping area between two target signals. Nuclear localization of TFEB was assessed by measuring the total nuclear intensity of TFEB, and the percentage of cells exceeding predefined nuclear intensity threshold was calculated. For Golgi fragmentation analysis, the abundance of GOLGA2- and TGOLN2-positive structures were quantified at the single-cell level. Cell-level data were exported for analysis using FlowJo software to develop contour plots and determine the percentage of cells showing fragmented Golgi morphologies.

#### Analysis of trans-Golgi network fragmentation

Golgi morphology was analyzed using the Morphology Explorer BioApplication. TGOLN2-postitive structures larger than 15 μm^2^ were classified as coalescent Golgi, whereas structures smaller than 15 μm^2^ were classified as fragmented Golgi. The number of fragmented Golgi per cell was quantified and compared between WT and *TRIM46* knockout cells.

#### Analysis of microtubule organization

Microtubule organization was quantified using the Cellomics Morphology Explorer Bioapplication. Microtubules were visualized by immunofluorescence staining with α-tubulin (Santa Cruz, sc-23948) as described above. The software classified microtubule structures as either spotted or fibrous based on morphology, and fibers smaller than 5 μm^2^ were defined as ‘short’ microtubules. To assess microtubule organization in *TRIM46* knockout cells, the ratio of the total area of short microtubules versus the total microtubule area was calculated and compared to WT. Microtubule bundling was analyzed using the built-in fiber alignment feature of the software. The orientation of individual fibers was determined relative to the image axis, and the standard deviation of fiber angles within a single cell was used as a measure of microtubule alignment. A minimum of 500 cells per well were analyzed, with data from multiple wells per experiment and ≥3 independent experiments pooled for quantitative analysis.

### Confocal microscopy and deconvolution

Sub-airy unit (0.6 AU) pinhole confocal microscopy with a Zeiss LSM900 or a Leica TCS-SP8 microscope was performed followed by computational image restoration with Huygens Essential (Scientific Volume Imaging) utilizing a constrained maximum likelihood estimation algorithm. Images were acquired using 63X/1.4NA plan apochromat oil immersion objective lenses and sampled at ideal Nyquist sampling rates in x, y, and z planes, allowing for sub-diffraction limited resolution following image restoration. All images were rendered on a high performance CUDA-GPA enabled workstation and 3D renders were generated for morphological analysis with Huygens Object Analyzer software. Sphericity values for LC3B-positive structures were obtained using the “RoughSphericity” feature. Colocalization between LC3B and TGOLN2 was quantified using the colocalization coefficients feature, reporting Pearson’s correlation coefficient. In addition, profile intensity plots were generated to visualize the overlap between LC3B and LAMP2 signals.

### Western blot analysis

Cells were washed twice with PBS and lysed in RIPA buffer (ThermoFisher, 89901) supplemented with protease inhibitors (Sigma,11836170001) and phosphatase inhibitors (Sigma, PHOSSTOP). Protein concentrations were determined using the BCA reagent (ThermoFisher, 23228). Equal amounts of protein were mixed with Laemmli Sample Buffer (Bio-Rad, 1610747) and boiled at 100°C for 10 min. Denatured proteins were separated on either 4–20% gradient or 10% SDS-PAGE gels (Bio-Rad) at 95 V and transferred to methanol-activated PVDF membranes or nitrocellulose membranes at 100 V for 1 h at 4°C. Membranes were blocked with 5% skim milk in PBS and incubated with primary antibodies overnight at 4°C. After three 10-minute washes with PBST (0.1% Tween-20), membranes were incubated with secondary antibodies for 1 h at room temperature. Following three additional washes with PBST, membranes were developed using ECL substrate (Bio-Rad, 1705061) for HRP-conjugated antibodies or directly imaged using a ChemiDoc imager (Bio-Rad) for fluorescent secondary antibodies. Protein quantification was performed using Image Lab software (Bio-Rad).

### Halo-LC3 assay for autophagy flux

HeLa WT or *TRIM46* knockout cells stably expressing Halo-LC3 (HT-LC3B) were seeded at a density of 3 × 10^5^ cells per well in 6-well plates. The following day, cells were pulsed with 2.5μM TMR-labeled HaloTag ligand (HL) for 30 min, followed by three washes with DMEM. Cells were then treated with either DMSO or 100 nM Bafilomycin A1 for 4 or 6 h. After treatment cells were washed twice with PBS and lysed in RIPA buffer supplemented with protease inhibitors. Protein concentrations were determined using a BCA assay, and equal amounts of protein were subjected to SDS-PAGE. Released HT/HL was detected by TMR fluorescence in-gel using a ChemiDoc™ MP Imager (Bio-Rad) prior to immunoblotting for actin.

### DQ-BSA assay

To assess lysosomal function, cells were seeded in a 96-well plate and incubated overnight with 10 µg/ml DQ-BSA (ThermoFisher, D12050) diluted in culture medium. Following incubation, cells were fixed with paraformaldehyde prior to immunolabeling with anti-LAMP2 and high content imaging.

### Membrane fractionation

Membrane fractionation was performed as described previously [69]. Cells from the two 15-cm dishes grown to confluence were harvested and homogenized in 5X cell pellet volume of buffer B1 buffer (20 mM HEPES-KOH, 400 mM sucrose, and 1 mM EDTA) supplemented with protease and phosphatase inhibitors and 0.3 mM DTT by passing through a 22-G needle. Homogenates were subjected to sequential differential centrifugation at 1000 x g for 10 min, 3000 × g for 10 min, 25, 000 × g for 20 min, and 100, 000 × g for 30 min to collect the pelleted membranes. Pellets were lysed in RIPA buffer supplemented with protease inhibitors and protein concentrations were determined using a BCA assay. Samples were mixed with Laemmli Sample Buffer, boiled for 10 min, and analyzed by SDS-PAGE and immunoblotting with antibodies against LC3B, LAMP2, and TGOLN2.

### Flow cytometric measurement of Golgiphagy

HeLa wild-type or *TRIM46* knockout cells (2 × 10^6^) were seeded in 10 cm dishes one day before transfection. Cells were transfected with 14 μg of the pKH116-Keima-YIPF3 plasmid (Addgene, 214970) using Lipofectamine 2000 (ThermoFisher) according to the manufacturer’s protocol. 16 h post-transfection, the culture medium was replaced with fresh medium. On day 2, cells were starved in Earle’s Balanced Salt Solution (EBSS; Thermo Fisher, 24010043) with or without 100 nM bafilomycin A1 for 6 h. After treatment, cells were harvested by trypsinization and resuspended in PBS containing 5% FBS for flow cytometry analysis. Acidified Golgi structures were assessed using dual-excitation ratiometric pH measurements at 405 nm (neutral pH ∼7) and 561 nm (acidic pH ∼4) laser excitations, with emission detected at 603/48 nm and 620/15 nm respectively using the flow cytometry (Attune NxT, Thermofisher). Flow cytometry data were acquired and analyzed using FlowJo software (version 10, Tree Star).

### Statistical analysis, graphing and figure assembly

All data are presented as mean ± s.e.m. Statistical significance between two groups was assessed using a two-sided unpaired Student’s *t*-test for normally distributed data or a two-sided Mann-Whitney *U*-test for non-normally distributed data. For multiple group comparison, one-way or two-way Anova followed by Tukey’s post hoc test was applied. Statistical analyses and graph generation were conducted using Graphpad Prism v10.6.1. Schematic diagrams were prepared with BioRender, and final figure assembly was performed using Adobe Illustrator.

## Acknowledgements

This work was supported by R01AI155746 to M.A.M from the US National Institutes of Health. S.O. was supported by T32AI007538 from the National Institute of Allergy and Infectious Disease. Confocal microscopy and flow cytometry were performed in the University of New Mexico Comprehensive Center shared resources, which are partially supported by P30CA118100 form the NIH. High content imaging was performed in the Autophagy, Inflammation, and Metabolism (AIM) Center core facility, which is supported by P20GM121176 from the NIH. Dr. Ruheena Javed (University of New Mexico) provided technical advice, and Drs. Thabata Duque, Tae-Hyung Kim, and Joseph Endicott (University of New Mexico) commented on the manuscript. Biorender software was used to generate graphics.

## Author contributions

S.O., S.U., B.S., and M.A.M performed research. S.O. and M.A.M. analyzed data and wrote the paper. The authors declare that they have no conflict of interest.

## Data availability statement

The data that support the findings of this study are available from the corresponding author, MAM, upon reasonable request,

## Abbreviations

AMPK: AMP-activated protein kinase
ATG3: autophagy related 3
ATG5: autophagy related 5
ATG7: autophagy related 7
ATG12: autophagy related 12
ATG13: autophagy related 13
ATG16L1: autophagy related 16 like 1
BECN1: beclin 1
CASM: conjugation of Atg8 to single membranes
GABARAP: GABA type A receptor-associated protein
GABARAPL1: GABA type A receptor associated protein like 1
GABARAPL2: GABA type A receptor associated protein like 2
GOLGA2: Golgin subfamily A member 2
HT: HaloTag
HL: HaloTag ligand
MAP1LC3A/LC3A: microtubule associated protein 1 light chain 3 alpha
MAP1LC3B/LC3B: microtubule associated protein 1 light chain 3 beta
MAP1LC3C/LC3C: microtubule associated protein 1 light chain 3 gamma
MTORC1: mechanistic target of rapamycin complex 1
PE: phosphatidyl ethanolamine
PIK3C3/VPS34: phosphatidylinositol 3-kinase catalytic subunit type 3
PS: phosphatidylserine
TECPR1: tectonin beta-propellor repeat containing 1
SQSTM1/p62: sequestosome 1
TFEB: transcription factor EB
TFE3: transcription factor binding to IGHM enhancer 3
TGOLN2: trans-golgi network protein 2
TRIM46: tripartite motif containing 46
ULK1: unc-51 like autophagy activating kinase 1
ULK2: unc-51 like autophagy activating kinase 2
VAIL: V-ATPase-ATG16L1 induced LC3 lipidation

