## Supplementary figures and table for "A role for CASM in the repair of damaged Golgi architecture"

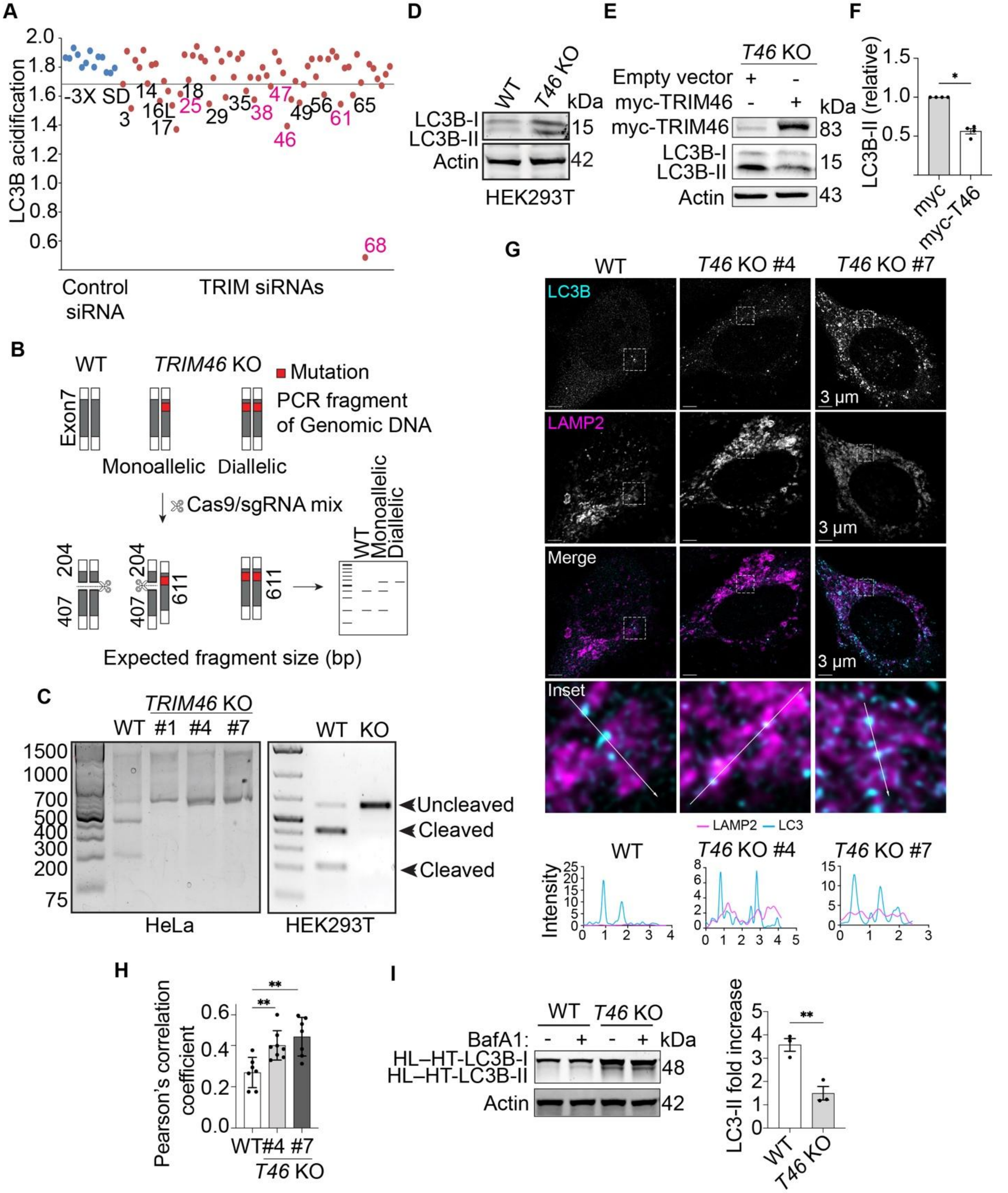


**Figure S1.** Confirmation of *TRIM46* knockout in HeLa and HEK293T cells. (**A**) A replicate of the experiment shown in Figure 1B. Numbers indicate TRIMs that were >3 standard deviations (line) below the mean measured for non-targeting siRNA (blue data points). Numbers in magenta were 'hits' in two out of two experiments. (**B**) Schematic diagram of the CRISPR confirmation assay. Genomic DNA was extracted, and PCR fragments containing the CRISPR/Cas9 target site in *TRIM46* were amplified and incubated with Cas9/sgRNA complexes. Cleavage of unedited DNA generates characteristic fragment sizes, but edited DNA lacks the cleavage site, thus enabling confirmation of CRISPR/Cas9-mediated editing. (**C**) Agarose gel images of HeLa (left) and HEK293T (right) cells confirming *TRIM46* knockout via Cas9/sgRNA cleavage of PCR fragments. In WT cells, intact *TRIM46* alleles were cleaved by Cas9/sgRNA, generating two fragments (204 bp and 407 bp). In *TRIM46* knockout cells, disruption of the cleavage site prevents further cutting, resulting in a single 611 bp fragment. (**D**) Immunoblot analysis of LC3B-II abundance in WT and *TRIM46* knockout (*T46* KO) HEK293T cells. (**E, F**) Immunoblot analysis of LC3B-II abundance in *TRIM46* knockout cells transfected with myc-TRIM46 or control vectors. Quantification of LC3B-II from 4 independent experiments. (**G, H**) Maximum image projection confocal micrographs of WT and *TRIM46* knockout HeLa cell lines stained with anti-LC3B and anti-LAMP2 antibodies. A zoomed-in image of the boxed region is shown below. Arrow designates the path of intensity profile measurements (plots, bottom). Plot (H) shows the Pearson's correlation coefficient from images in G. Each point represents data from a different confocal image. (**I**) WT and *TRIM46* knockout cells expressing Halo-LC3B were pulsed with TMR-labeled HL for 3 min prior to a 6-hour chase in full media in the presence or absence of BafA1. HL–HT-LC3B was detected by in-gel fluorescence and actin detected by immunoblot. Plot shows the abundance of HL–HT-LC3B-II in BafA1-treated cells relative to that in cells treated with DMSO vehicle only. Statistical analyses were performed using an unpaired Student t-test (F, I) or ordinary one-way ANOVA followed by Dunnett's multiple-comparison test (H). Data: mean ± SEM; *, $p < 0.05$; **, $p < 0.01$.

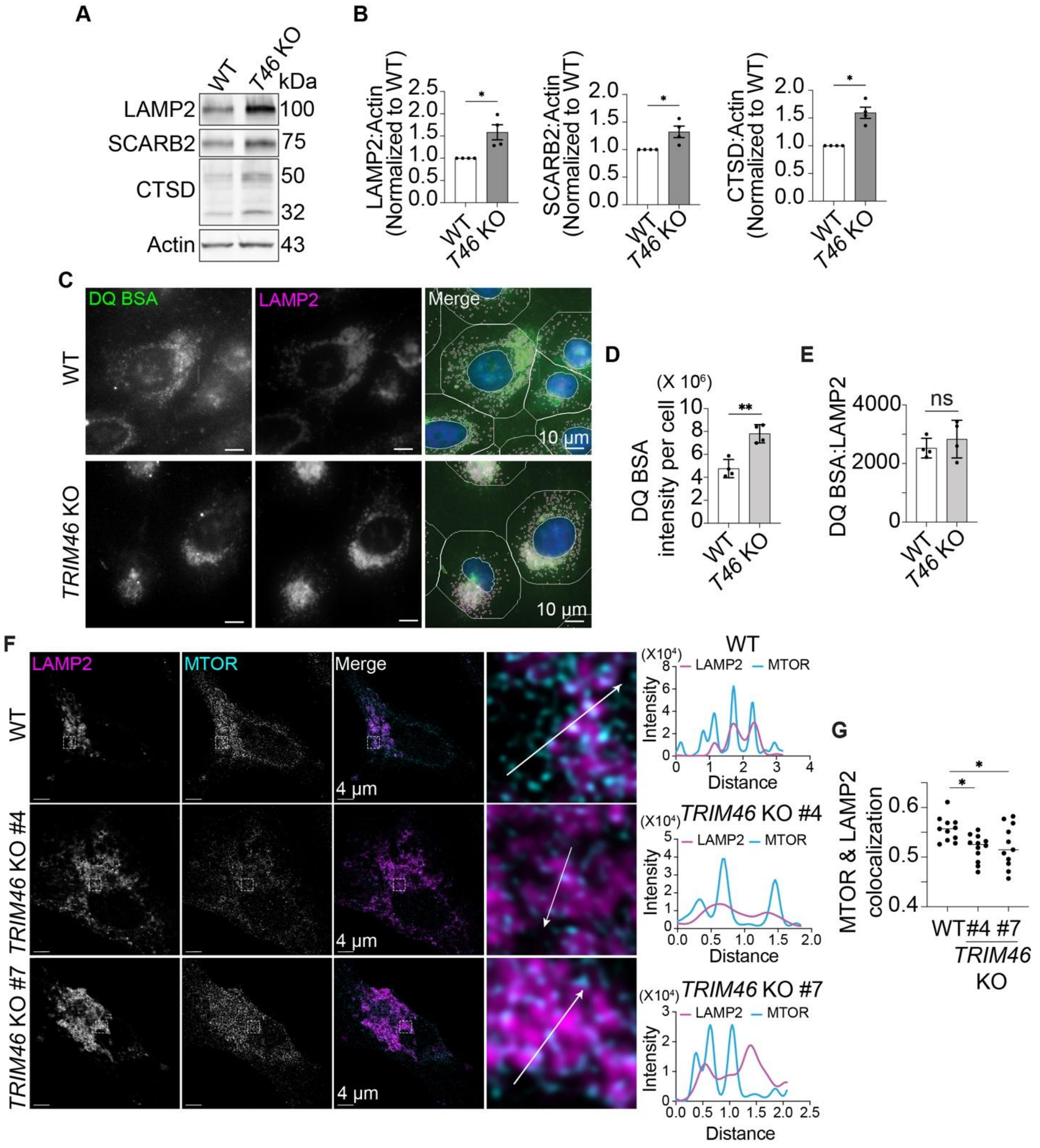


**Figure S2.** Lysosomal biogenesis is increased in *TRIM46* knockout cells. (**A, B**) Immunoblot analysis of lysosomal protein abundance in WT and *TRIM46* knockout HEK293T cells. Plots (**B**) show quantification of the indicated protein's abundance relative to WT. Each data point represents an independent experiment. (**C-E**) DQ-BSA assay for lysosomal function. WT and *TRIM46* knockout HeLa cells were treated with 10 µg/mL DQ-BSA for 6 h prior to fixation and high-content imaging. Plots show the DQ-BSA intensity per cell (D), and ratio of DQ-BSA intensity to LAMP2 area per cell (E). Each data point represents the average calculated from >500 cells. (**F**) Maximum image projections of confocal images showing LAMP2 and MTOR staining in WT and *TRIM46* knockout HeLa cells. The area in the box is enlarged and shown in the microscopy image to the right. Arrow shows the track of the intensity profile measurements (plots, right). (**G**) Colocalization between MTOR and LAMP2 analyzed using Pearson's correlation coefficient. Each point represents data from a different confocal image. Statistical analyses were performed using an unpaired Student t-test (B, D, E) or ordinary one-way ANOVA followed by Dunnett's multiple-comparison test (G). Data: mean ± SEM; *, $p < 0.05$; **, $p < 0.01$; ns, not significant.

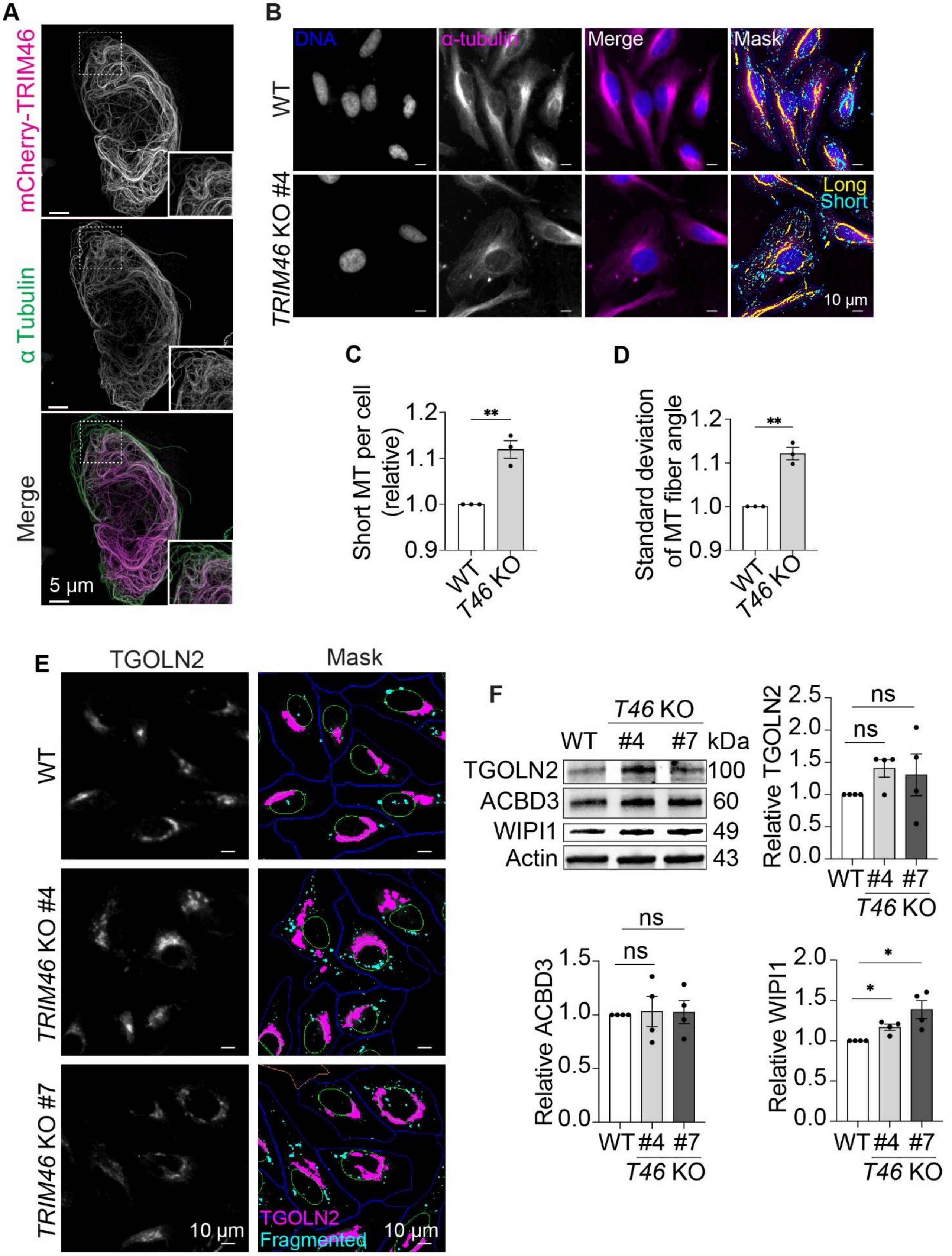

A
mCherry-TRIM46
α Tubulin
Merge
5 μm
B
DNA
α-tubulin
Merge
Mask
WT
TRIM46 KO #4
Long
Short
10 μm
C
Short MT per cell (relative)
1.2
1.1
1.0
0.9
**
WT
T46 KO
D
Standard deviation of MT fiber angle
1.2
1.1
1.0
0.9
**
WT
T46 KO
E
TGOLN2
Mask
WT
TRIM46 KO #4
TRIM46 KO #7
10 μm
TGOLN2
Fragmented
10 μm
F
T46 KO
WT
#4
#7
kDa
TGOLN2
100
ACBD3
60
WIPI1
49
Actin
43
Relative TGOLN2
2.5
2.0
1.5
1.0
0.5
0.0
ns
ns
WT #4 #7
T46 KO
Relative ACBD3
1.5
1.0
0.5
0.0
ns
ns
WT #4 #7
T46 KO
Relative WIPI1
2.0
1.5
1.0
0.5
0.0
*
*
WT #4 #7
T46 KO

**Figure S3.** TRIM46 localizes to microtubules and regulates microtubule organization and Golgi fragmentation. (**A**) Maximum image projection confocal images of HeLa cells transfected with mCherry-TRIM46 and stained for α-tubulin. (**B-D**) High-content imaging analysis of microtubule organization in WT and *TRIM46* knockout (shown, KO #4) HeLa cells. WT and *TRIM46* knockout HeLa cells were fixed and stained with anti-α-tubulin antibodies. (B) Short (α-tubulin structures < 5 $\mu m^2$; cyan mask) and long (α-tubulin structures > 5 $\mu m^2$; yellow mask) microtubule structures were automatically segmented. (C) The relative abundance of short microtubules per cell was quantified as a percentage of total microtubules and compared between WT and of *TRIM46* knockout cells. (D) The standard deviation of angles of long microtubule fibers, a measure of microtubule disorganization, was calculated per cell, and the mean values were compared between WT and *TRIM46* knockout cells. Data points represent three independent experiments, each based on an average of >500 cells. (**E**) Representative high-content microscopy images of WT and *TRIM46* knockout HeLa cells stained with TGOLN2 related to Figure 3B and 3C. Magenta mask illustrates “large” TGOLN2 structures (>15 $\mu m^2$) and cyan mask highlights fragmented TGOLN2 structures (<15 $\mu m^2$). (**F**) Immunoblot analysis of the abundance of selected Golgi-resident proteins in WT and *TRIM46* knockout HeLa cells. Plots show quantification of relative protein levels in *TRIM46* knockout cells compared to WT. Dots represents independent experiments. Statistical analyses were performed using an unpaired Student t-test (C, D) or non-parametric Mann-Whitney U test (F). Data: mean ± SEM; *, $p < 0.05$; **, $p < 0.01$; ns, not significant.

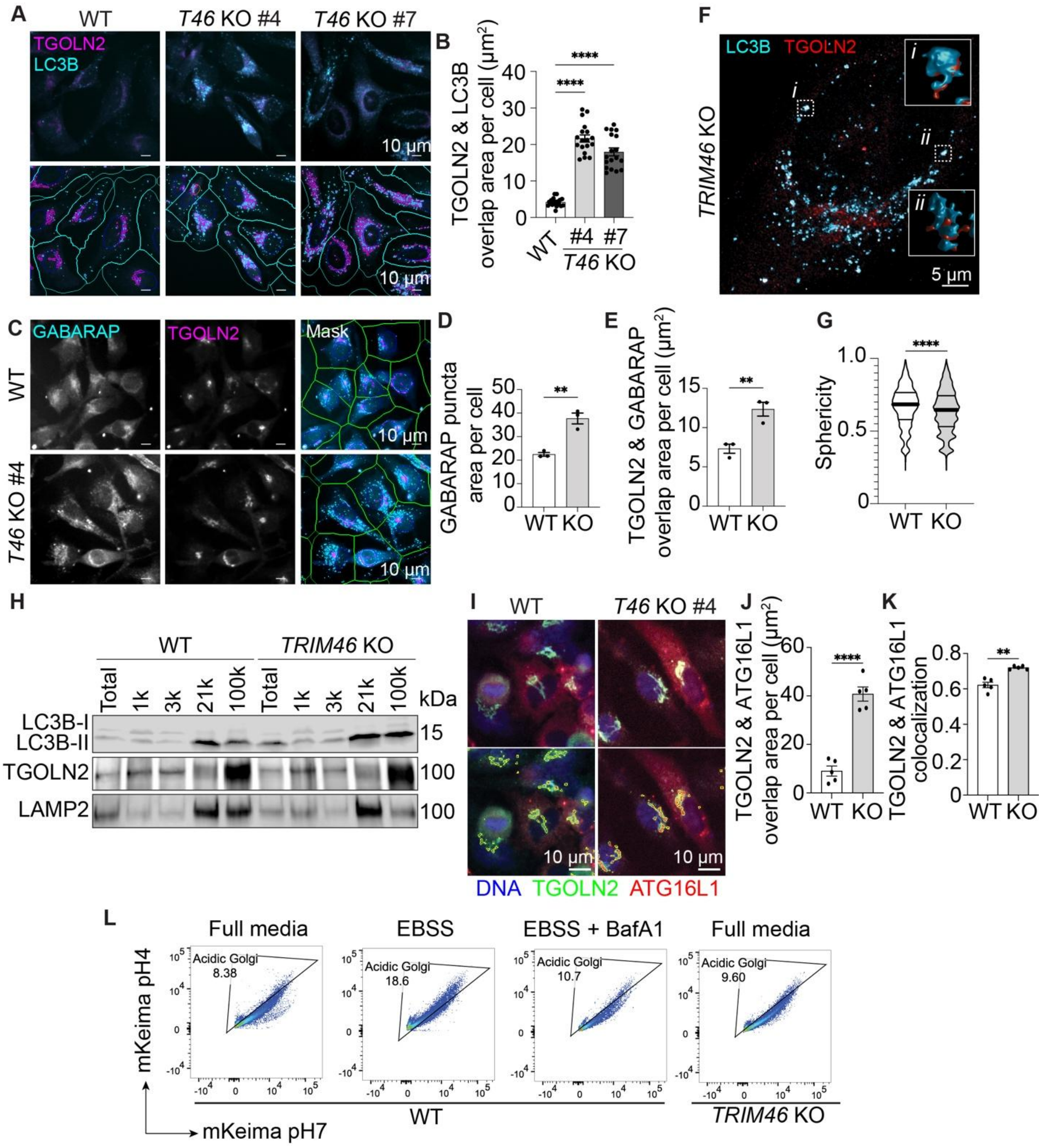


**Figure S4.** *TRIM46* knockout cells show increased non-degradative Atg8ylation of the trans-Golgi network. (**A, B**) High content imaging analysis of colocalized LC3B and TGOLN2 in WT and two *TRIM46* knockout clones. (**C-E**) High content image analysis of WT and *TRIM46* knockout HeLa cells stained with antibodies against GABARAP and TGOLN2. The average abundance of GABARAP puncta, and GABARAP/TGOLN2 overlap area were quantified per cell. Each data point represents the average of >500 cells. (**F, G**) Confocal microscopy analysis of the morphology of structures showing overlapping TGOLN2 and LC3B signal. Maximum image projection of a *TRIM46* knockout HeLa cell. Three dimensional reconstructions of the boxed structures are shown on the right. Violin plot (G) showing the sphericity of LC3B-positive structures quantitated from confocal images of WT and *TRIM46* knockout HeLa cells. Thick bars, median; thin bars, quartiles. Perfectly spherical objects will have a value of 1. (**H**) Gradient centrifugation of membrane fractions isolated from WT and *TRIM46* knockout cells. (**I-K**) High content imaging analysis of TGOLN2 and ATG16L1 colocalization in WT or *TRIM46* knockout HeLa cells. Cells were transfected with mCherry-ATG16L1 and stained with TGOLN2. Masks in bottom image: yellow, TGOLN2 positive; red, ATG16L1 positive. Graphs show the abundance of overlapping signal (J) and the Pearson's correlation (K) on a per-cell basis. Each data point represents the average of >500 cells. (**L**) Flow cytometric analysis of the delivery of the Golgiphagy reporter YIPF3-mKeima to lysosomes in WT and *TRIM46* knockout cells. WT HeLa cells were starved in EBSS for 6 h with or without 100 nM BafA1 as controls. One representative experiment of two was shown. Statistical analyses were performed using an ordinary one-way ANOVA followed by Dunnett's multiple-comparison test (B) or unpaired Student t-test (D, E, G, J, K). Data: mean ± SEM; **, $p < 0.01$; ****, $p < 0.0001$.

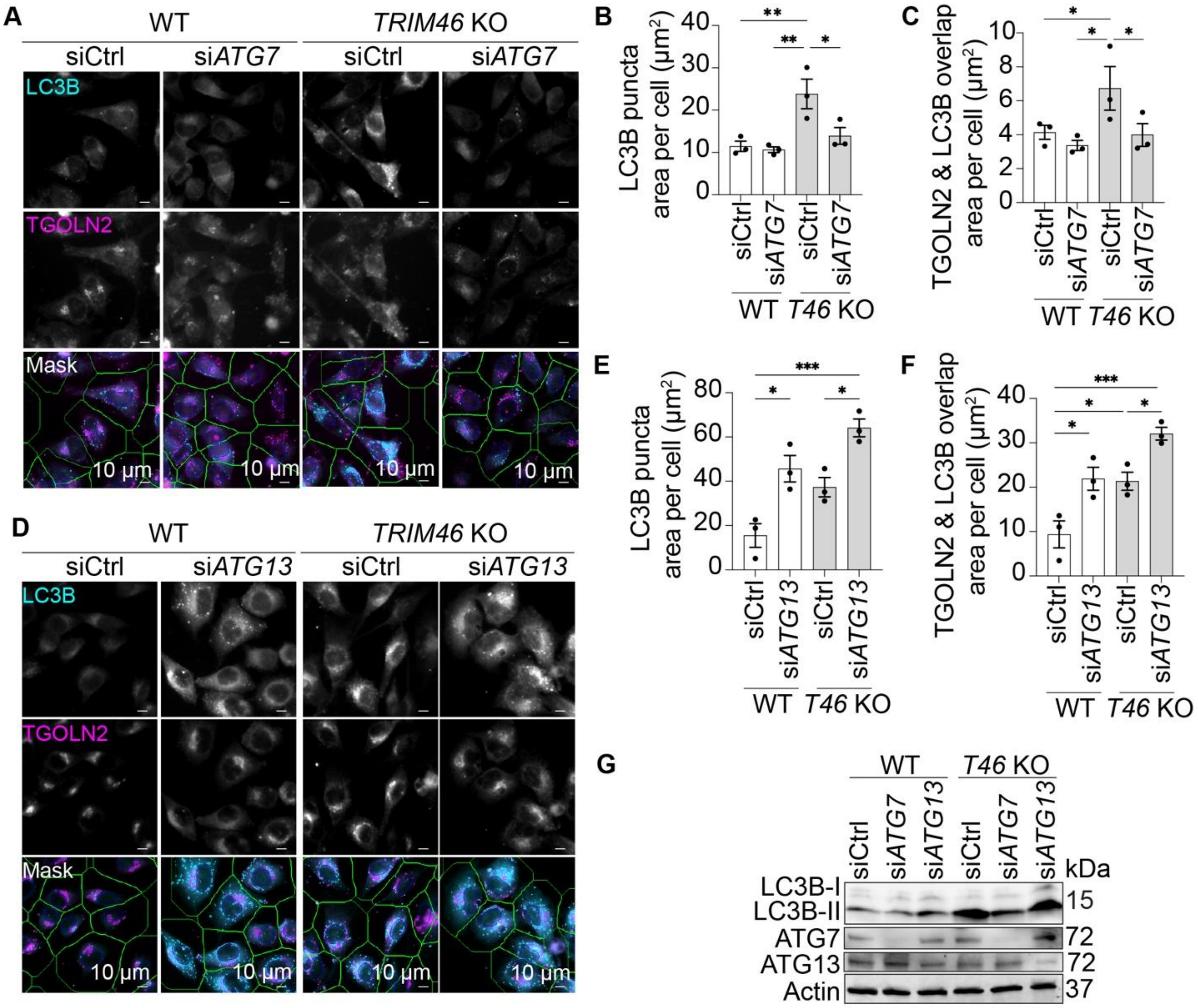


**Figure S5.** ATG7, but not ATG13, is required for Atg8ylation in *TRIM46* knockout cells. (**A, D**) Representative images of WT and *TRIM46* knockout HeLa cells transfected with siRNAs targeting *ATG7* (A) or *ATG13* (D), stained with antibodies against LC3B and TGOLN2. Plots show quantification of LC3B puncta area per cell (**B, E**) and colocalized LC3B and TGOLN2 signal (**C, F**). Data points represent biological replicates, each based on an average of more than 500 cells. (**G**) Western blot confirming ATG7 or ATG13 knockdown efficiency. Statistical analyses were performed using two-way ANOVA followed by Tukey's multiple-comparison test. Data: mean ± SEM; *, $p < 0.05$; **, $p < 0.01$; ***, $p < 0.001$.

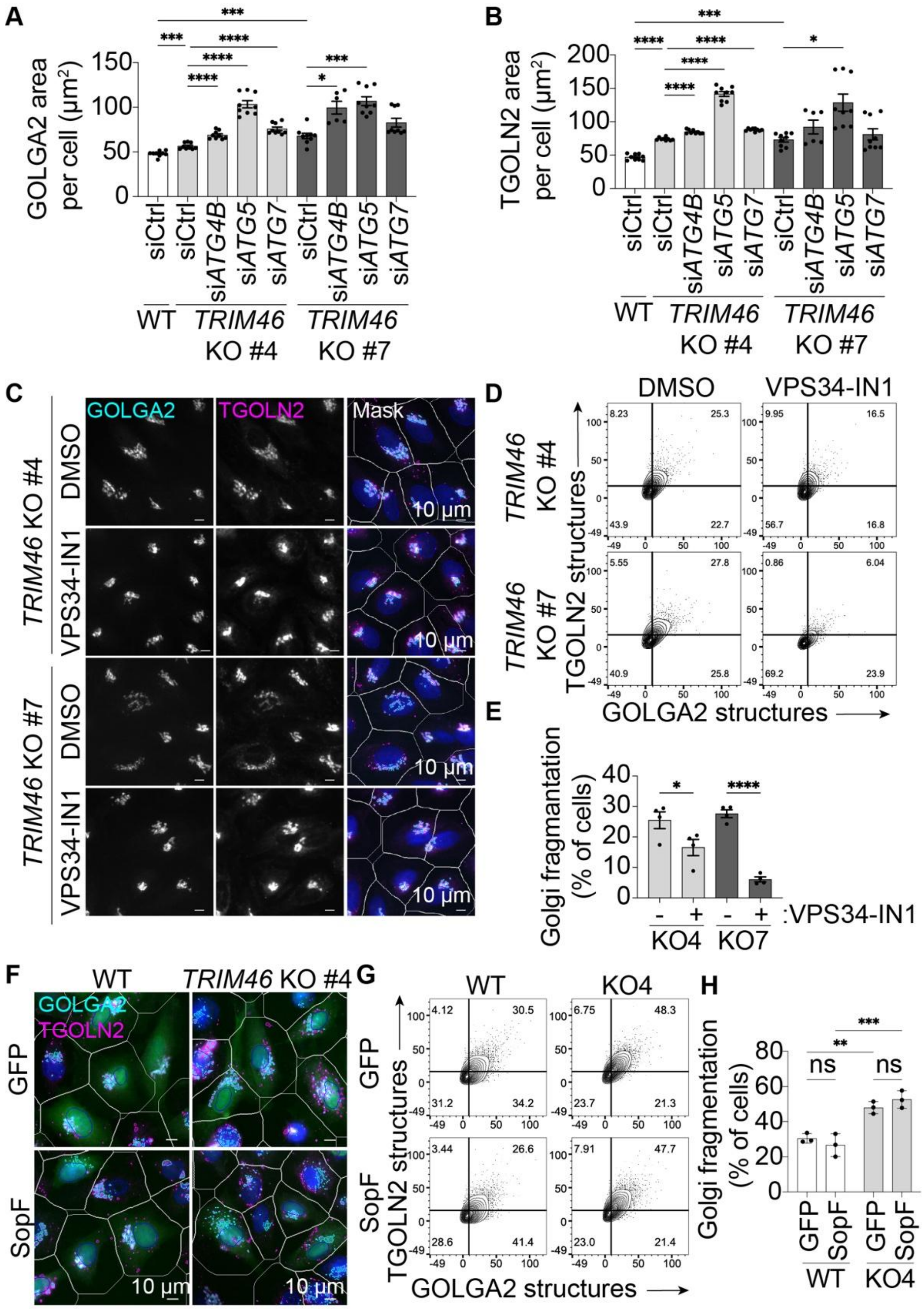


**Figure S6.** Impacts of CASM or VPS34 inhibition on Golgi architecture in *TRIM46* knockout cells. (**A-B**) Quantification of GOLGA2 and TGOLN2 area per cell, corresponding to Fig. 5B. Data points represent biological replicates; each calculated from the average of >500 cells. (**C-E**) *TRIM46* knockout HeLa cells were treated with DMSO or 10 μM of VPS34-IN1 for 4 h prior to immunostaining. (C) Representative high content images of two clones of *TRIM46* knockout HeLa cells stained with antibodies against GOLGA2 and TGOLN2. (D) Contour plots show the number of distinct GOLGA2 and TGOLN2 structures per cell. The numbers in the upper right-hand quadrants indicate the percentage of cells exhibiting Golgi fragmentation, defined as >10 GOLGA2$^{+}$ structures and >15 TGOLN2 $^{+}$ structures per cell. N ≥3394 cells analyzed per sample. (E) Percentage of cells classified as exhibiting fragmented Golgi structures. (**F**) Representative high content images of WT or *TRIM46* knockout HeLa cells transfected with either GFP or SopF (co-expressing GFP) and stained for GOLGA2 and TGOLN2. (**G**) Contour plots showing the number of distinct GOLGA2- and TGOLN2-positive structures per cell. N ≥ 9000 cells were analyzed per sample. (**H**) Plot showing the percentage of cells classified as exhibiting fragmented Golgi structures. Statistical analyses were performed using Welch's ANOVA followed by Dunnett's multiple-comparison test (A, B) or two-way ANOVA followed by Tukey's multiple-comparison test (E, H). Data: mean ± SEM; *, $p < 0.05$; **, $p < 0.01$; ***, $p < 0.001$; ****, $p < 0.0001$; ns, not significant.

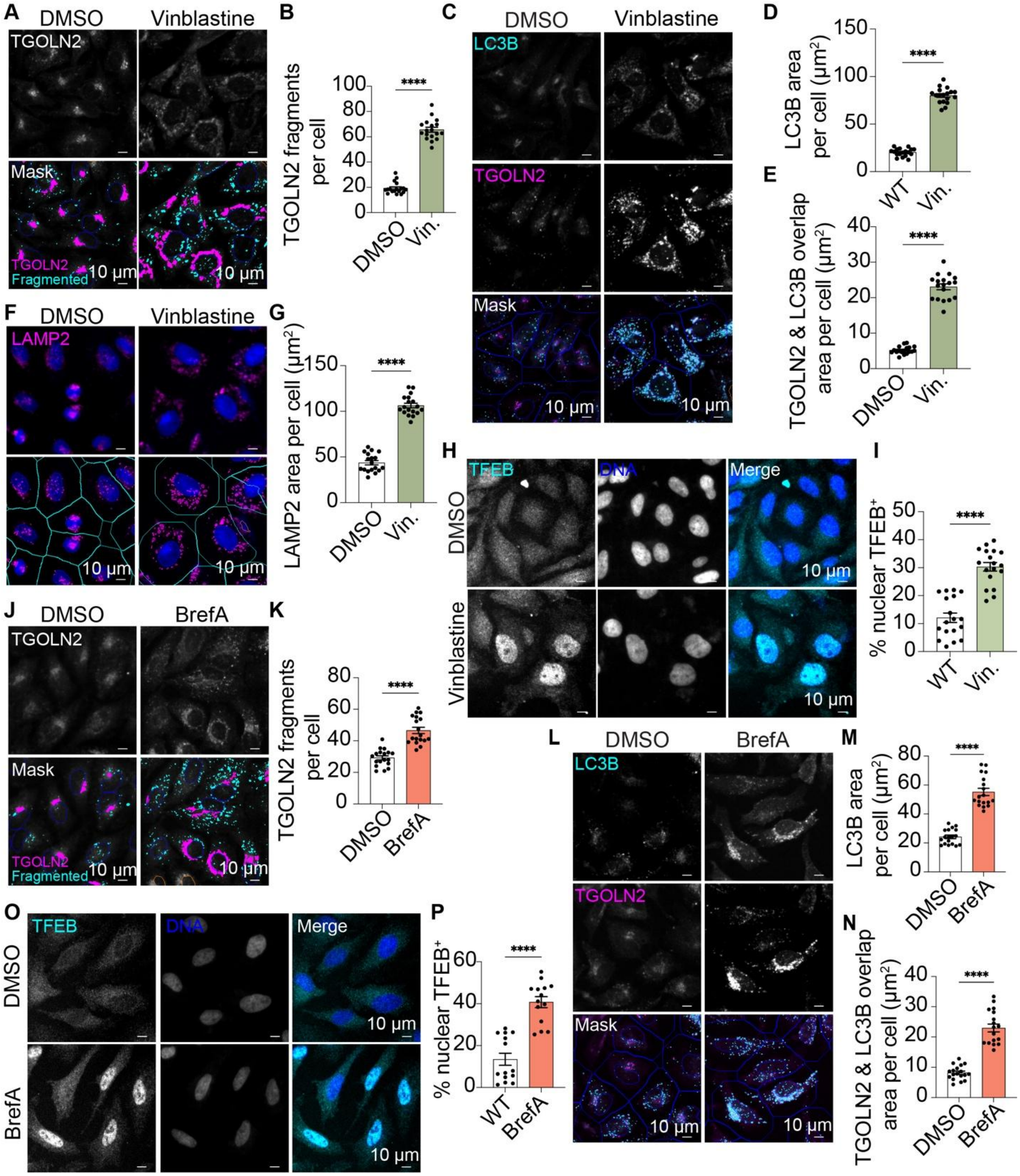


**Figure S7.** Microtubule disruption with vinblastine triggers Atg8ylation of Golgi fragments and increased lysosomal biogenesis. WT HeLa cells were treated with 100 nM vinblastine or DMSO control for 16 h. (**A-B**) High-content imaging analysis of TGOLN2 fragmentation measured by counting small TGOLN2-positive structures (cyan mask) smaller than 15 μm². Larger TGOLN2-positive structures are labeled with the magenta mask. (**C-E**) Representative high-content microscopy images of TGOLN2 (magenta) and LC3B (cyan) colocalization following vinblastine treatment. Plots show quantitation of LC3B puncta area per cell (D) and overlapping area between TGOLN2 and LC3B (E). (**F, G**) High content imaging analysis of WT HeLa cells measuring LAMP2 abundance following vinblastine or DMSO control treatment. (**H, I**) High content imaging analysis of TFEB nuclear localization following vinblastine or control treatment of HeLa cells. (**J-P**) High content image analysis of Golgi fragmentation (J, K), LC3B puncta abundance (L, M), TGOLN2 colocalization with LC3B (L, N), and TFEB nuclear localization (O, P) in HeLa cells treated or not with brefeldin A (BrefA) for 16 h. Each data point represents the average of more than 500 cells. Statistical analyses were performed using an unpaired Student t-test. Data: mean ± SEM; ****, $p < 0.0001$.

**Table 1.** Reagents.

| REAGENT or RESOURCE | SOURCE | IDENTIFIER |
|---|---|---|
| **Antibodies** | | |
| Anti-mCherry antibody | Abcam | ab183628, RRID: AB_2650480 |
| GABARAP | Abcam | ab109364, RRID: AB_10861928 |
| NPC2 | Abcam | ab218192, RRID: AB_2941808 |
| Purified Mouse Anti-SQSTM1/p62 Ick ligand | BD biosciences | P0067, RRID: AB_398151 |
| Goat Anti-Mouse IgG (H+L)-HRP | Bio-Rad Laboratories | 1721011, RRID: AB_2617113 |
| Goat Anti-Rabbit IgG (H+L)-HRP | Bio-Rad Laboratories | 1721019, RRID: AB_11125143 |
| TGOLN2/TGN46 | Bio-Rad Laboratories | AHP500G, RRID: AB_2203291 |
| PRKAA/AMPK | Cell Signaling Technology | 2532, RRID: AB_330331 |
| phospho-PRKAA/AMPK T172 | Cell Signaling Technology | 2535, RRID: AB_331250 |
| ATG13 (E1Y9V) Rabbit mAb | Cell Signaling Technology | 13468, RRID: AB_2797419 |
| BECN1 | Cell Signaling Technology | 3459, RRID: AB_560924 |
| GOLGA2/GM130 | Cell Signaling Technology | 12480, RRID: AB_2797933 |
| phospho-RPS6KB/p70 S6K | Cell Signaling Technology | 9205, RRID: AB_330944 |
| TFEB | Cell Signaling Technology | 4240, RRID: AB_11220225 |
| MTOR | Cell Signaling Technology | 2983, RRID: AB_2105622 |
| ULK1 | Cell Signaling Technology | 8054, RRID: AB_11178668 |
| IRDye® 680LT Goat anti-Mouse IgG Secondary Antibody | LI-COR Biosciences | 925-68020, RRID: AB_2687826 |
| IRDye 800CW Goat anti-Mouse IgG Secondary Antibody | LI-COR Biosciences | 925-32210, RRID: AB_2687825 |

| | | |
|---|---|---|
| LC3B | MBL | PM036, RRID: AB_2274121 |
| Anti-Halotag monoclonal antibody | Promega | G921A, RRID:AB_2688011 |
| CTSD/cathepsin D | Proteintech | 21327-1-AP, RRID: AB_10733646 |
| SCARB2/LIMP2 | Proteintech | 27102-1-AP, RRID: AB_2880756 |
| TRIM46 | Proteintech | 21026-1-AP, RRID: AB_10732843 |
| Cas9 | Santa Cruz Biotechnology | sc-517386, RRID: AB_2800509 |
| LAMP2 | Santa Cruz Biotechnology | sc-18822, RRID: AB_626858 |
| RPS6KB/p70 S6K | Santa Cruz Biotechnology | sc-8418, RRID: AB_628094 |
| Anti-ACTB/actin Antibody (2Q1055) | Santa Cruz Biotechnology | sc-58673, RRID: AB_2223345 |
| TUBA/alpha-tubulin | Santa Cruz Biotechnology | sc-23948, RRID: AB_628410 |
| Rabbit Anti-LC3B | Sigma Aldrich | L7543, RRID: AB_796155 |
| ANTI-FLAG | Sigma Aldrich | F1804, RRID: AB_262044 |
| ACBD3/GCP60 | Thermo Fisher Scientific | MA5-25999, RRID: AB_2723827 |
| TGOLN2 | Thermo Fisher Scientific | MA5-37930, RRID: AB_2897850 |
| WIPI1 | Thermo Fisher Scientific | PA5-34973, RRID: AB_2552322 |
| HCS CellMask™ Near-IR Stain | Thermo Fisher Scientific | H32722 |
| Goat anti-Rabbit IgG (H+L) Highly Cross-Adsorbed Secondary Antibody, Alexa Fluor 488 | Thermo Fisher Scientific | A-11034, RRID: AB_2576217 |
| Goat anti-Mouse IgG (H+L) Highly Cross-Adsorbed Secondary Antibody, Alexa Fluor Plus 647 | Thermo Fisher Scientific | A32728, RRID: AB_2633277 |
| **Bacterial and virus strains** | | |
| NEB 5-alpha Competent *E. coli* (High Efficiency) | New England Biolabs | C2987 |
| XL10-Gold Ultracompetent cells | Agilent Technologies | 210518 |
| **Chemicals, peptides, and recombinant protein** | | |

| | | |
|---|---|---|
| 10x Tris/Glycine/SDS buffer | Bio-Rad Laboratories | 1610732 |
| 2x Laemmli Buffer | Bio-Rad Laboratories | 1610737 |
| 4x Laemmli Buffer | Bio-Rad Laboratories | 1610747 |
| Clarity ECL | Bio-Rad Laboratories | 1705061 |
| Glycine | Bio-Rad Laboratories | 1610718 |
| Tris base | Bio-Rad Laboratories | 1610719 |
| Vinblastine | Cayman chemical | 11762 |
| Vps34-IN1 | Cayman chemical | 17392 |
| Bovine serum albumin | Fisher Scientific | CAS 9048-46-8 |
| DTT (Dithiothreitol) | Gold Biotechnology | DTT10 |
| Bafilomycin $A_1$ | Invivogen | tlrl-baf1 |
| HaloTag® TMR Ligand | Promega | G8251 |
| Brefeldin A | Sigma Aldrich | B6542 |
| Ethylenediaminetetraacetic acid disodium salt dihydrate | Sigma Aldrich | E5134 |
| 2-mercaptoethanol | Sigma Aldrich | M3148 |
| Monensin | Sigma Aldrich | M5273 |
| Nocodazole | Sigma Aldrich | M1404 |
| PHOSSTOP | Sigma Aldrich | 4906837001 |
| cOmplete™, Mini, EDTA-free Protease Inhibitor Cocktail | Sigma Aldrich | 11836170001 |
| Puromycin dihydrochloride | Sigma Aldrich | P9620 |
| Saponin | Sigma Aldrich | 84510 |
| Sucrose | Sigma Aldrich | 84097 |
| Tween 20 | Sigma Aldrich | P1379 |
| Millipore® Steriflip® Vacuum Tube Top Filter | Sigma Aldrich | SE1M003M00 |
| Dulbecco's modified Eagle's medium | Thermo Fisher Scientific | 11965084 |
| Earle's Balanced Salt Solution | Thermo Fisher Scientific | 24010043 |
| fetal bovine serum | Thermo Fisher Scientific | 26140095 |
| Hoechst 33342 | Thermo Fisher Scientific | H3570 |
| Lipofectamine 2000 Reagent | Thermo Fisher Scientific | 11668019 |

| | | |
|---|---|---|
| Opti-MEM Reduced Serum Medium | Thermo Fisher Scientific | 31985070 |
| penicillin-streptomycin | Thermo Fisher Scientific | 15140122 |
| Restore Plus Western Blot Stripping Buffer | Thermo Fisher Scientific | 46430 |
| RIPA lysis buffer | Thermo Fisher Scientific | 89901 |
| Trypsin-EDTA (0.25%), phenol red | Thermo Fisher Scientific | 25200072 |
| Ampicillin sodium salt | VWR | IC19014805 |
| Dimethyl sulfoxide (DMSO) | VWR | EMMX14586 |
| Ethanol | VWR | 89125172 |
| Kanamycin sulfate | VWR | 97061-600 |
| Methanol | VWR | BDH20291GLP |
| Paraformaldehyde | VWR | JTS8987 |
| Potassium chloride | VWR | EMPX14051 |
| Potassium phosphate monobasic | VWR | EMDPX15651 |
| 2-Propanol (isopropyl Alcohol) | VWR | BDH20271GLP |
| Sodium chloride | VWR | BDH928625KG |
| Sodium phosphate dibasic | VWR | 97061472 |
| Triton X-100 | VWR | EM9410 |
| **Critical commercial assays** | | |
| Agilent QuikChange Lightning Site-Directed Mutagenesis Kit | Agilent Technologies | 210518 |
| BCA reagent | Thermo Fisher Scientific | 23228 |
| Extracta Plus DNA | QuantaBio | 95213-050 |
| Gateway™ BP Clonase™ II Enzyme mix | Thermo Fisher Scientific | 11789100 |
| Gateway™ LR Clonase™ II Enzyme mix | Thermo Fisher Scientific | 11791020 |
| Guide-it™ Complete sgRNA Screening System | Takara Bio | 632636 |
| QIAprep Spin Miniprep Kit | Qiagen | 27104 |
| QIAquick PCR Purification Kit | Qiagen | 28106 |

| | | |
|---|---|---|
| PureLink™ HiPure Plasmid Midiprep Kit | Thermo Fisher | K210005 |
| ProFection Mammalian Transfection System | Promega | E1200 |
| **Experimental models: Cell lines** | | |
| HeLa *TRIM46* knockout | This study | |
| HEK293T *TRIM46* knockout | This study | |
| HeLa stably expressing HT-LC3 | This study | |
| HeLa *TRIM46* knockout cells stably expressing HT-LC3 | This study | |
| **Oligonucleotides** | | |
| *TRIM46* Forward primer for sgRNA | This study | 5'-CCTCTAATACGACTCACTATAGGTACCGTTGAGTTCCGGCGCAGTTTAAGAGCTATGC-3' |
| *TRIM46* Forward primer for genomic DNA amplification | This study | 5'-GCTGCTTTCCCTTTTCCT-3' |
| *TRIM46* Reverse primer for genomic DNA amplification | This study | 5'-CTCTGAAGTTCAGAGAGGGT-3' |
| *TRIM46* Forward primer for site-directed mutagenesis g1407t | This study | 5'-AGTTCCGGCGCACTGATGTGCCTGCTC-3' |
| *TRIM46* Reverse primer for site-directed mutagenesis g1407t | This study | 5'-GAGCAGGCACATCAGTGCGCCGGAACT-3' |
| siControl | GE Dharmacon | D-001206-13-05 |
| si*ATG4B* | GE Dharmacon | M-005786-01-0005 |
| siATG5 | GE Dharmacon | M-004374-04-0005 |
| siATG7 | GE Dharmacon | M-020112-01-005 |
| siATG13 | GE Dharmacon | M-020765-01-0005 |
| siBECN1 | GE Dharmacon | M-010552-01-0005 |
| siULK1 | GE Dharmacon | M-005049-00-0005 |
| si*MID2/TRIM1* | GE Dharmacon | M-007076-01 |
| siTRIM2 | GE Dharmacon | M-006955-00 |

| | | |
|---|---|---|
| siTRIM3 | GE Dharmacon | M-006931-00 |
| siTRIM4 | GE Dharmacon | M-007101-00 |
| siTRIM5 | GE Dharmacon | M-007100-00 |
| siTRIM6 | GE Dharmacon | M-007121-01 |
| siTRIM7 | GE Dharmacon | M-007077-01 |
| siTRIM10 | GE Dharmacon | M-006920-01 |
| siTRIM11 | GE Dharmacon | M-007075-00 |
| siTRIM13 | GE Dharmacon | M-006923-00 |
| siTRIM14 | GE Dharmacon | M-010976-00 |
| siTRIM15 | GE Dharmacon | M-007102-01 |
| siTRIM16 | GE Dharmacon | M-012220-01 |
| siTRIM16L | GE Dharmacon | M-023055-01 |
| siTRIM17 | GE Dharmacon | M-006981-01 |
| si*MID1/TRIM18* | GE Dharmacon | M-006537-01 |
| si*PML/TRIM19* | GE Dharmacon | M-006547-01 |
| si*MEFV/TRIM20* | GE Dharmacon | M-011081-00 |
| siTRIM21 | GE Dharmacon | M-006563-02 |
| siTRIM22 | GE Dharmacon | M-006927-03 |
| siTRIM23 | GE Dharmacon | M-006523-00 |
| siTRIM24 | GE Dharmacon | M-005387-03 |
| siTRIM25 | GE Dharmacon | M-006585-00 |
| siTRIM26/ZNF173 | GE Dharmacon | M-019558-02 |
| siTRIM27 | GE Dharmacon | M-006552-01 |
| siTRIM28 | GE Dharmacon | M-005046-01 |
| siTRIM29 | GE Dharmacon | M-012409-01 |
| siTRIM31 | GE Dharmacon | M-006939-01 |
| siTRIM32 | GE Dharmacon | M-006950-01 |
| siTRIM33 | GE Dharmacon | M-005392-03 |
| siTRIM34 | GE Dharmacon | M-006997-01 |
| siTRIM35 | GE Dharmacon | M-006952-02 |
| siTRIM37 | GE Dharmacon | M-006538-02 |
| siTRIM38 | GE Dharmacon | M-006929-01 |

| | | |
|---|---|---|
| siTRIM39 | GE Dharmacon | M-007028-01 |
| siTRIM40 | GE Dharmacon | M-007129-01 |
| siTRIM41 | GE Dharmacon | M-007105-02 |
| siTRIM42 | GE Dharmacon | M-007173-00 |
| siTRIM43 | GE Dharmacon | M-007127-01 |
| siTRIM44 | GE Dharmacon | M-017337-01 |
| siTRIM45 | GE Dharmacon | M-007073-01 |
| siTRIM46 | GE Dharmacon | M-007071-01 |
| siTRIM47 | GE Dharmacon | M-007106-02 |
| siTRIM48 | GE Dharmacon | M-007059-01 |
| siTRIM49 | GE Dharmacon | M-007030-01 |
| siTRIM50 | GE Dharmacon | M-007130-00 |
| siTRIM51 | GE Dharmacon | M-010079-02 |
| siTRIM52 | GE Dharmacon | M-007095-00 |
| siTRIM54 | GE Dharmacon | M-007032-01 |
| siTRIM55 | GE Dharmacon | M-007092-01 |
| siTRIM56 | GE Dharmacon | M-007079-00 |
| siTRIM58 | GE Dharmacon | M-013985-02 |
| siTRIM59 | GE Dharmacon | M-007172-01 |
| siTRIM60 | GE Dharmacon | M-007153-00 |
| siTRIM61 | GE Dharmacon | M-028281-01 |
| siTRIM62 | GE Dharmacon | M-007010-02 |
| siTRIM63 | GE Dharmacon | M-007093-01 |
| siTRIM64 | GE Dharmacon | M-026740-04 |
| siTRIM65 | GE Dharmacon | M-018490-01 |
| siTRIM66 | GE Dharmacon | M-026772-01 |
| siTRIM67 | GE Dharmacon | M-032288-01 |
| siTRIM68 | GE Dharmacon | M-007007-01 |
| siTRIM71 | GE Dharmacon | M-023459-01 |
| siTRIM72 | GE Dharmacon | M-032293-02 |
| siTRIM73 | GE Dharmacon | M-028896-01 |
| siTRIM74 | GE Dharmacon | M-031736-01 |

| | | |
|---|---|---|
| si*CMYA5/TRIM76* | GE Dharmacon | M-016373-01 |
| **Recombinant DNA** | | |
| mCherry | Pankiv et al, 2007 [1] | |
| mCherry-ATG16L1 | Kumar et al, 2021 [2] | |
| pDONR221-hTRIM46 | DNASU | HsCD00862242 |
| mCherry-TRIM46 | This study | N/A |
| MYC-TRIM46 | This study | N/A |
| mKeima-YIPF3 | Addgene | 214970 |
| pLV[CRISPR]-hCas9:T2A:Puro-U6-hTRIM46 | VectorBuilder | VB900138-6978huh |
| pLEX307-Halo LC3 | Javed et al, 2025 [3] | |
| pRP-EGFP-CMV-Flag-SopF | Vectorbuilder | VB251216-1400bmp |
| **Software and algorithms** | | |
| Prism 8 | GraphPad | N/A |
| Image Lab | BIO-RAD LABORATORIES | N/A |
| FlowJo (v10.10.0) | BD Biosciences | N/A |
| iDEV software | Thermo Fisher Scientific | N/A |
| Huygens Object Analyzer and Colocalization | Scientific Volume Imaging | N/A |
| LASX acquisition software | Leica | N/A |
| BioRender | BioRender.com | N/A |
| ICE software | Synthego | |


[1] Pankiv S, Clausen TH, Lamark T, et al. p62/SQSTM1 binds directly to Atg8/LC3 to facilitate degradation of ubiquitinated protein aggregates by autophagy. The Journal of biological chemistry. 2007;282(33):24131-45.

[2] Kumar S, Javed R, Mudd M, et al. Mammalian hybrid pre-autophagosomal structure HyPAS generates autophagosomes. Cell. 2021;184(24):5950-5969 e22.

[3] Javed R, Jain A, Duque T, et al. Mammalian ATG8 proteins maintain autophagosomal membrane integrity through ESCRTs. The EMBO journal. 2023;42(14):e112845.
